# Structural basis for PRC2 decoding of active histone methylation marks H3K36me2/3

**DOI:** 10.1101/2020.04.22.054684

**Authors:** Ksenia Finogenova, Jacques Bonnet, Simon Poepsel, Ingmar B. Schäfer, Katja Finkl, Katharina Schmid, Claudia Litz, Mike Strauss, Christian Benda, Jürg Müller

## Abstract

Repression of genes by Polycomb requires that PRC2 modifies their chromatin by trimethylating lysine 27 on histone H3 (H3K27me3). At transcriptionally active genes, di- and trimethylated H3K36 inhibit PRC2. Here, the cryo-EM structure of PRC2 on dinucleosomes reveals how binding of its catalytic subunit EZH2 to nucleosomal DNA orients the H3 N-terminus via an extended network of interactions to place H3K27 into the active site. Unmodified H3K36 occupies a critical position in the EZH2-DNA interface. Mutation of H3K36 to arginine or alanine inhibits H3K27 methylation by PRC2 on nucleosomes *in vitro*. Accordingly, *Drosophila* H3K36A and H3K36R mutants show reduced levels of H3K27me3 and defective Polycomb repression of HOX genes. The relay of interactions between EZH2, the nucleosomal DNA and the H3 N-terminus therefore creates the geometry that permits allosteric inhibition of PRC2 by methylated H3K36 in transcriptionally active chromatin.

## INTRODUCTION

Many post-translational modifications on histone proteins are essential for processes in the underlying chromatin. Typically, histone modifications themselves do not alter chromatin structure directly but function by binding effector proteins which alter chromatin or by interfering with such interactions. The histone methyltransferase Polycomb Repressive Complex 2 (PRC2) and its regulation by accessory proteins and histone modifications represent a prime example for understanding these interaction mechanisms (Laugesen et al., 2019; Yu et al., 2019). PRC2 trimethylates lysine 27 in histone H3 (H3K27me3), a modification that is essential for the transcriptional repression of developmental regulator genes that control cell fate decisions in metazoans (McKay et al., 2015; Pengelly et al., 2013). H3K27me3 marks chromatin for interaction with PRC1, an effector which compacts chromatin (Francis et al., 2004; Grau et al., 2011). H3K27me3 is also recognized by PRC2 itself, and this interaction allosterically activates the PRC2 enzyme complex to facilitate deposition of H3K27me3 across extended domains of chromatin (Hansen et al., 2008; Jiao and Liu, 2015; Margueron et al., 2009). Genetic studies and subsequent biochemical work established that PRC2 is in addition subject to negative regulation. In particular, the H3K4me3, H3K36me2 and H3K36me3 marks present on nucleosomes in transcriptionally active chromatin directly inhibit H3K27 methylation by PRC2 (Gaydos et al., 2012; Klymenko and Müller, 2004; Schmitges et al., 2011; Streubel et al., 2018; Yuan et al., 2011). Importantly, while stimulation of PRC2 activity by H3K27me3 acts *in trans*, inhibition of PRC2 by H3K4me3, H3K36me2 and H3K36me3 requires that these modifications are present *in cis*, that is, on the same H3 molecule containing the K27 substrate lysine (Schmitges et al., 2011; Yuan et al., 2011). While recent structural studies have uncovered the allosteric activation mechanism for PRC2 (Jiao and Liu, 2015; Justin et al., 2016), the molecular basis of PRC2 inhibition by active chromatin marks has remained enigmatic. In particular, in nucleosome binding assays, PRC2-DNA interactions make the largest contribution to the nucleosome-binding affinity of PRC2 (Choi et al., 2017; Wang et al., 2017) and H3K4me3, H3K36me2 and H3K36me3 do not seem to have a major effect on this binding affinity (Guidotti et al., 2019; Jani et al., 2019; Schmitges et al., 2011). Instead, these three modification were found to reduce the k_cat_ of PRC2 for H3K27 methylation (Jani et al., 2019; Schmitges et al., 2011). Recent cross-linking studies led to the suggestion of a possible sensing pocket for H3K36 on the surface of EZH2 (Jani et al., 2019) but there is no structural data how that proposed interaction might occur. Similarly, a recent structure of PRC2 bound to a dinucleosome revealed how the catalytic lobe of PRC2 contacts nucleosomes through DNA interactions but provided no structural insight into how the H3 N-termini might be recognized (Poepsel et al., 2018). Here, a refined structure of PRC2 bound to a dinucleosome allowed us to visualize how the histone H3 N-terminus on substrate nucleosomes is threaded into the EZH2 active site. Our analyses reveal that H3K36 assumes a critical position in the PRC2-nucleosome interaction interface that permits the complex to gauge the H3K36 methylation state.

## RESULTS

### EZH2 interaction with nucleosomal DNA orients the H3 N-terminus for H3K27 binding to the active site

We assembled recombinant full-length human PRC2 in complex with its accessory factor PHF1 (i.e. PHF1-PRC2) (Choi et al., 2017) on a heterodimeric dinucleosome (di-Nuc), which consisted of a ‘substrate’ nucleosome with unmodified histone H3 and an ‘allosteric’ nucleosome containing H3 with a trimethyllysine analog (Simon et al., 2007) at K27, separated by a 35 base pair (bp) DNA linker (Poepsel et al., 2018) (**Figure 1A, B**). Single particle cryo-electron microscopy analysis yielded a reconstruction of the PHF1-PRC2:di-Nuc assembly with an overall resolution of 5.2 Å (**Supplementary Figures 1–3**). The map showed clear density for the catalytic lobe of PRC2 with similar chromatin interactions and binding geometry as previously described for the catalytic lobe of AEBP2-PRC2 (Poepsel et al., 2018) where PRC2 contacts the two nucleosomes via interactions with the DNA gyres (**Figure 1C**). Specifically, the substrate nucleosome is bound by the EZH2_CXC_ domain residues K563, Q565, K569 and Q570 (**Figure 1D, Supplementary Figure 4A**, cf. (Poepsel et al., 2018)), while the allosteric nucleosome is contacted by EED and by the SBD and SANT1 domains of EZH2 (**Figure 1E**, cf. (Poepsel et al., 2018)). We could not detect density for the ‘bottom lobe’ of PRC2 (Chen et al., 2018; Kasinath et al., 2018) or for the N-terminal winged-helix and tudor domains of PHF1 that bind DNA and H3K36me3, respectively (Ballaré et al., 2012; Cai et al., 2013; Choi et al., 2017; Li et al., 2017; Musselman et al., 2013).

**FIGURE 1.**
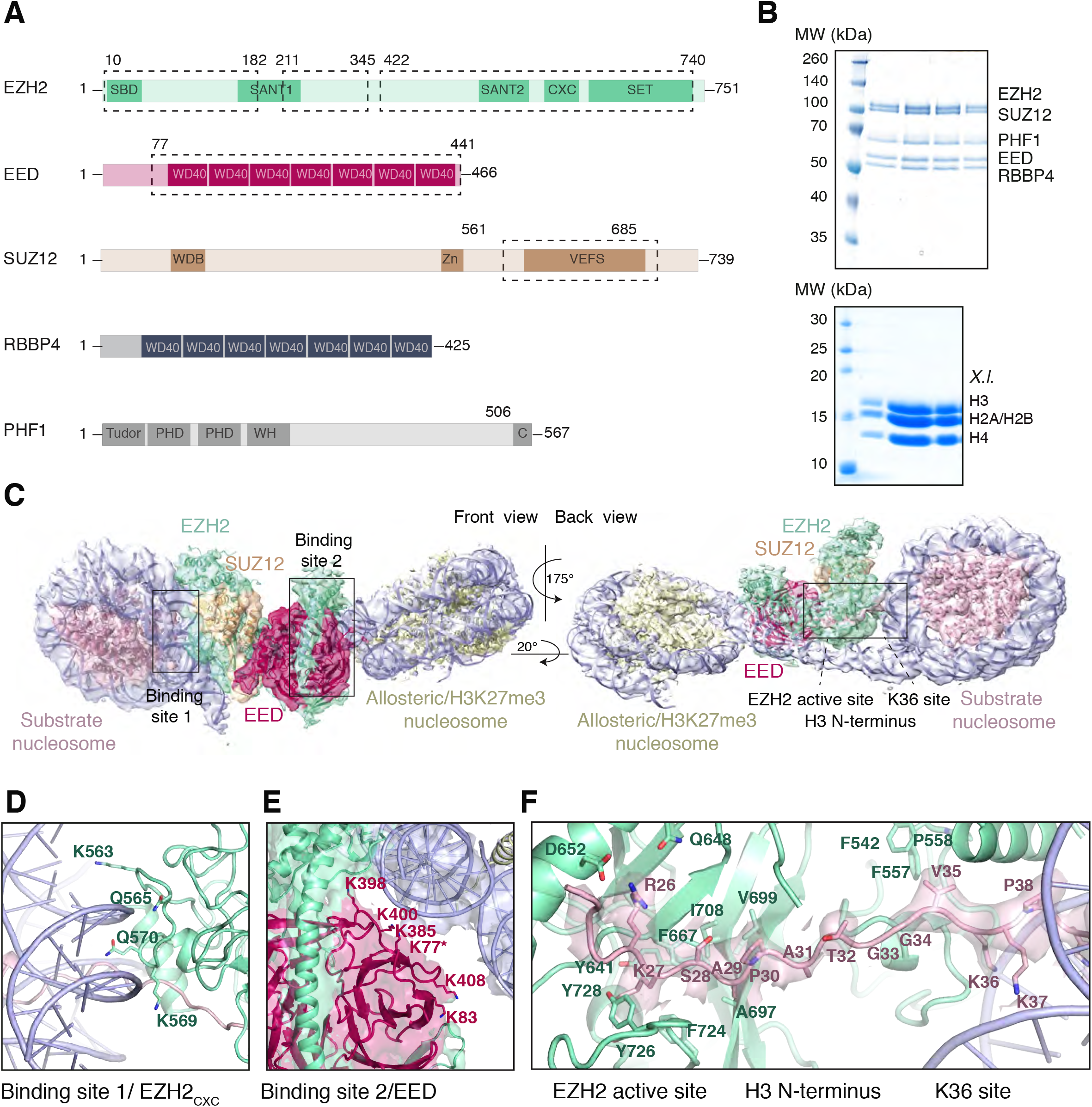
Interaction of the PRC2 catalytic lobe with nucleosomal DNA orients the H3 N-terminus for H3K27 binding to the active site. (**A**) Domain organization in the five subunits of PHF1-PRC2. Dashed boxes indicate protein portions visible in the PHF1-PRC2:di-Nuc cryo-EM reconstruction and fitted in the structural model. In PHF1, C corresponds to the short C-terminal fragment used in PHF1_C_-PRC2. (**B**) Coomassie-stained SDS PAGE analysis of representative PHF1-PRC2 (upper panel) and *X.l*. octamer preparations (lower panel) after size-exclusion chromatography (SEC) purification. Pooled fractions of PHF1-PRC2, incubated with heterodimeric dinucleosomes generated by DNA ligation of a reconstituted unmodified and a H3Kc27me3-modified mononucleosome were used as input material for cryo-EM analysis. (**C**) Cryo-EM reconstruction of PHF1-PRC2:di-Nuc in two orientations with fitted crystal structures of human PRC2 catalytic lobe (PDB: 5HYN, (Justin et al., 2016)) and nucleosomes (1AOI, (Luger et al., 1997)) in a di-Nuc model with 35 bp linker DNA (see also **Supplementary Figures 1–4, Table S1, Movie S1**). Density is colored as in (**A**) to show PRC2 subunits, DNA (blue) and octamers of substrate (pink) and allosteric (yellow) nucleosomes. Boxes indicate regions shown in (**D**), (**E**) and (**F**), respectively. (**D**) Interaction of EZH2_CXC_ residues with the DNA gyres of the substrate nucleosome; residues mutated in PRC2^CXC>A^ are indicated. For the H3 N-terminus (pink), only the peptide backbone is shown in this view (see **F**). (**E**) Interface formed by EED and the EZH2 SBD domain with DNA gyres on the allosteric nucleosome; residues mutated in PRC2^EED>A^ are indicated. Asterisk indicates the approximate location of a residue, which is not built in the model. (**F**) The H3 N-terminus (pink), shown as a pseudoatomic model fitted into the 4.4 Å density map, is recognized by EZH2 through an extensive interaction network (see text). Note the well-defined side-chain density of H3K36 (see also **Supplementary Figure 3D and 4C-E**).

Using particle signal subtraction and focused refinement on the interface of EZH2 and the substrate nucleosome (**Supplementary Figures 2–3**), we then obtained an improved map at an apparent overall resolution of 4.4 Å which revealed well-defined density for the H3 N-terminus (**Figure 1F, Supplementary Figure 3B-D**). The visible sidechain density combined with the crystallographic models of the PRC2 catalytic lobe and of the mononucleosome enabled us to build a pseudo-atomic model of the histone H3 N-terminus spanning residues R26 to K37 (**Figure 1F**). This model revealed that EZH2 recognizes the H3 N-terminus via an extended network of contacts besides the previously described ionic interactions near the active site where H3 R26 interacts with EZH2 Q648/D652, and H3 K27 with the aromatic cage above the EZH2 catalytic center (Justin et al., 2016) (**Figure 1F**). Specifically, our structure suggests two hydrophobic hotspots, the first one involving H3 A29/P30 and EZH2 residues F667, A697, V699, I708 and F724 and the second one involving H3 V35 and F542, F557 and P558 of EZH2 (**Figure 1F**). H3 G33/G34 is likely not recognized by PRC2 but might act as a flexible hinge between the two hydrophobic interaction sites (**Figure 1F**). H3K36 is directly juxtaposed to the EZH2_CXC_-DNA interaction surface and appears to be involved in the EHZ2-DNA interface. The side chain density of H3K36 suggests that the epsilon-amino group of H3K36 engages in a polar interaction with the carbonyl group of Q570 and in long-range electrostatic interactions with the phosphate backbone of the nucleosomal DNA (**Figure 1F, Supplementary Figure 4 C-E**). Taken together, our analyses reveal that an extensive network of interactions between EZH2, the nucleosomal DNA and the H3 N-terminus. This complex geometric arrangement orients the H3 N-terminus into an extended conformation, threading H3K27 into the EZH2 active site. In this context, it should be noted that a previously postulated H3K36-binding pocket centered on E579 of EZH2 (Jani et al., 2019) is located approximately 19 Å away from H3K36 in our structure (**Supplementary Figure 4F**). An interaction of H3K36 with E579 of EZH2 as proposed by Muir and co-workers (Jani et al., 2019) would require a very different binding geometry of PRC2 on the nucleosome and major structural rearrangements of PRC2 or the nucleosome in order to avoid steric clashes.

### The EZH2 CXC contact with DNA is essential for H3K27 methylation

We next analysed how the PRC2 surfaces contacting the substrate and the allosteric nucleosome contribute to the formation of productive PRC2-chromatin interactions. For these experiments, we used PHF1_C_-PRC2, which contains the minimal 5-kDa PRC2-interaction domain of PHF1 (**Figure 1A**, (Chen et al., 2020; Choi et al., 2017)) but lacks the H3K36me3-binding tudor and the DNA-binding winged-helix domains of PHF1 (Choi et al., 2017; Li et al., 2017; Musselman et al., 2013). PHF1_C_-PRC2 therefore only retains the DNA-binding surfaces of the 4-subunit PRC2 core complex and was used because it generally behaved better in purifications than the 4-subunit PRC2 core complex. For simplicity we shall, in the following, refer to the PHF1_C_-PRC2 complex as PRC2. We generated three mutant versions of PRC2. In PRC2^CXC>A^ (K563A Q565A K569A Q570A), the EZH2_CXC_ interface is mutated (**Figure 1D**), in PRC2^EED>A^ (K77A K83A K385A K398A K400A K408A), the EED interface contacting the allosteric nucleosome (**Figure 1E**), is mutated, and PRC2^CXC>A/EED>A^ carries the combination of these mutations. We first used electromobility shift assays (EMSA) to measure the binding affinity of wild-type and mutant PRC2 complexes on mononucleosomes. These mononucleosomes were assembled on a 215 bp long DNA fragment containing the 147-bp 601 nucleosome-positioning sequence (Lowary and Widom, 1998) in the center and linker DNA on both sides. Wild-type PRC2 bound this mononucleosome with an apparent Kd in the mid-nanomolar range (**Figure 2A, B**, cf. (Choi et al., 2017)). The binding affinities of PRC2^CXC>A^ or PRC2^EED>A^ were two- to three-fold lower than that of wild-type PRC2 and that of PRC2^CXC>A/EED>A^ was about five-fold lower compared to the wild-type complex (**Figure 2A, B**, compare lanes 11-30 with 1-10, and **Supplementary Figure 5A**). The residual nucleosome binding shown by PRC2^CXC>A/EED>A^ (**Figure 2A**, lanes 21-30) could in part be due to incomplete disruption of the mutated interfaces but in part is likely also due to the previously identified nucleosome-binding activity of the PRC2 bottom lobe (Chen et al., 2018; Nekrasov et al., 2005). The overall binding affinity of the PRC2 core complex for chromatin therefore appears to result from interactions of at least three distinct surfaces of this complex with nucleosomes. Among those, binding of the EZH2_CXC_ domain to the nucleosomal DNA contributes only modestly to the total binding affinity.

**FIGURE 2.**
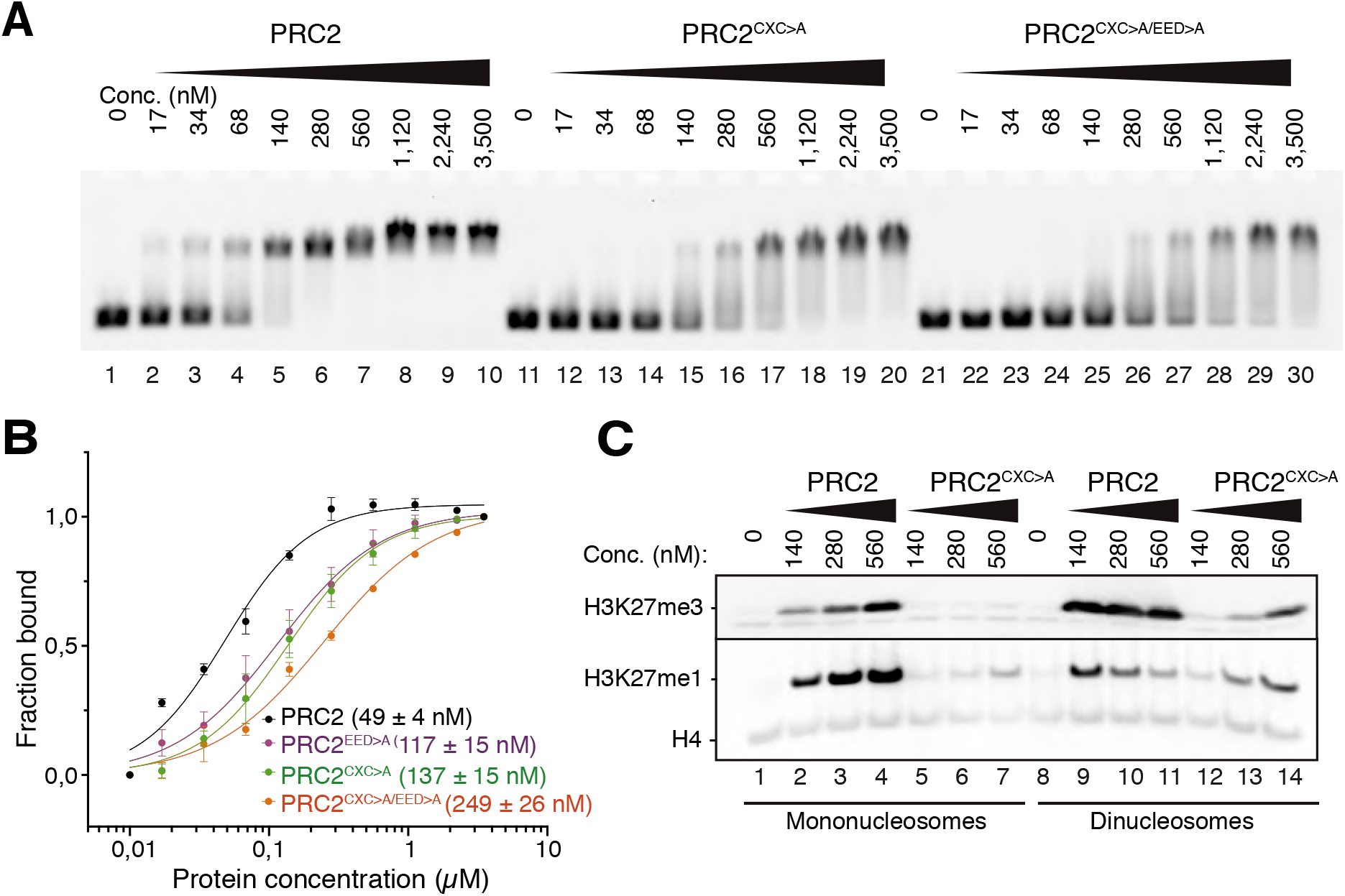
The EZH2_CXC_-DNA interaction interface is critical for H3K27 methylation on nucleosomes. (**A**) Binding reactions with indicated concentrations of PRC2 (lanes 1-10), PRC2^CXC>A^ (lanes 11-20) or PRC2^CXC>A/EED>A^ (lanes 21-30) and 45 nM 6-carboxyfluorescein-labeled mononucleosomes, analyzed by EMSA on 1.2% agarose gels; for analysis of PRC2^EED>A^ binding, see **Supplementary Figure 5A**. (**B**) Quantitative analysis of EMSA data in A by densitometry of 6-carboxyfluorescein signals from independent experiments (n=3); error bars, SEM. (**C**) Western Blot (WB) analysis of H3K27me1 and H3K27me3 formation in HMTase reactions with indicated concentrations of PRC2 and PRC2^CXC>A^ on 446 nM mononucleosomes (lanes 1-7) or 223 nM dinucleosomes (lanes 8-14). Note that these concentrations result in equal numbers of nucleosomes and therefore equal numbers of H3 substrate molecules in the reactions on mono- and dinucleosomes, as can be seen from the Coomassie-stained gel of the reactions in **Supplementary Figure 5B**. H4 WB signal served as control for Western blot processing.

We then analysed the histone methyltransferase (HMTase) activity of PRC2^CXC>A^. On the same mononucleosomes used above, PRC2^CXC>A^ showed almost no detectable HMTase activity compared to wild-type PRC2 (**Figure 2C**, compare lanes 5-7 with 2-4, see also **Supplementary Figure 5B**). On dinucleosomes, EED binding to one nucleosome might be expected to facilitate interaction of the mutated EZH2^CXC>A^ domain with the H3 N-termini on the juxtaposed second nucleosome. Indeed, on dinucleosomes, the PRC2^CXC>A^ complex does generate H3K27me1 and -me3 but less efficiently than wild-type PRC2 (**Figure 2C**, compare lanes 12-14 with 9-11). When comparing the activities of the different complexes, it should be kept in mind that the interpretation of H3K27me3 formation as read-out of complex activity on dinucleosomes is more complicated than on mononucleosomes, because H3K27me3, once placed on one of the nucleosomes, will allosterically activate PRC2 to methylate H3K27 on the linked second nucleosome (Margueron et al., 2009) (Jiao and Liu, 2015). The main conclusion to be drawn from our analyses here is that the DNA-binding interaction of the EZH2_CXC_ domain with substrate nucleosomes is critical for engaging the H3 N-terminus in a manner that allows H3K27 methylation.

### Unmodified H3K36 in the EZH2_CXC_-DNA interaction interface is critical for H3K27 methylation in nucleosomes

The architecture of the EZH2_CXC_-DNA interface around H3K36 (**Figure 1F**) suggested that a bulkier side chain, such as that of a tri- or di-methylated lysine or an arginine may not be accommodated in this interface. In EMSAs, the affinity of PRC2 for binding to mononucleosomes containing a trimethyllysine analog at H3K36 (H3Kc36me3) was indistinguishable from that for binding to unmodified mononucleosomes (**Figure 3A, B**). However, as previously reported (Schmitges et al., 2011; Yuan et al., 2011), H3K27 mono- and trimethylation by PRC2 was strongly inhibited on H3Kc36me3-containing mononucleosomes (**Figure 3C**, compare lanes 5-7 with 2-4, see also **Supplementary Figure 6A**). Methylation of H3K27 was also inhibited on mononucleosomes where H3K36 had been mutated to arginine or alanine (H3^K36R^ and H3^K36A^ mononucleosomes, respectively) (**Figure 3C**, compare lanes 8-13 with 2-4). PRC2 inhibition on H3^K36R^ and H3^K36A^ mononucleosomes was, however, less severe than on H3Kc36me3 mononucleosomes (**Figure 3C**, compare lanes 8-13 with 5-7). We note that the quantitative analyses here show inhibition of PRC2 HMTase activity on H3^K36A^ mononucleosomes, consistent with earlier studies (Jani et al., 2019), whereas other studies previously had failed to detect inhibition on H3^K36A^ mononucleosomes (Schmitges et al., 2011). Taken together, our structural and biochemical analyses suggest that the unmodified side chain of H3K36 is critical for the productive positioning of H3K27 in the catalytic center of PRC2. Neither the bulkier side chains of trimethyllysine or arginine nor the short apolar side chain of alanine appear to provide the correct fit at the position of H3K36. Finally, we compared PRC2 HMTase activity on histone H3_18-42_ peptides that were either unmodified or contained H3K36me3 using a Masspectrometry-based methylation assay. Importantly, on this isolated peptide, H3K36me3 did not inhibit H3K27 monomethylation by PRC2 (**Figure 3D, Supplementary Figure 6B**). The allosteric inhibition of PRC2 by H3K36me3 therefore only occurs in the context of the geometric constraints of the nucleosome.

**FIGURE 3.**
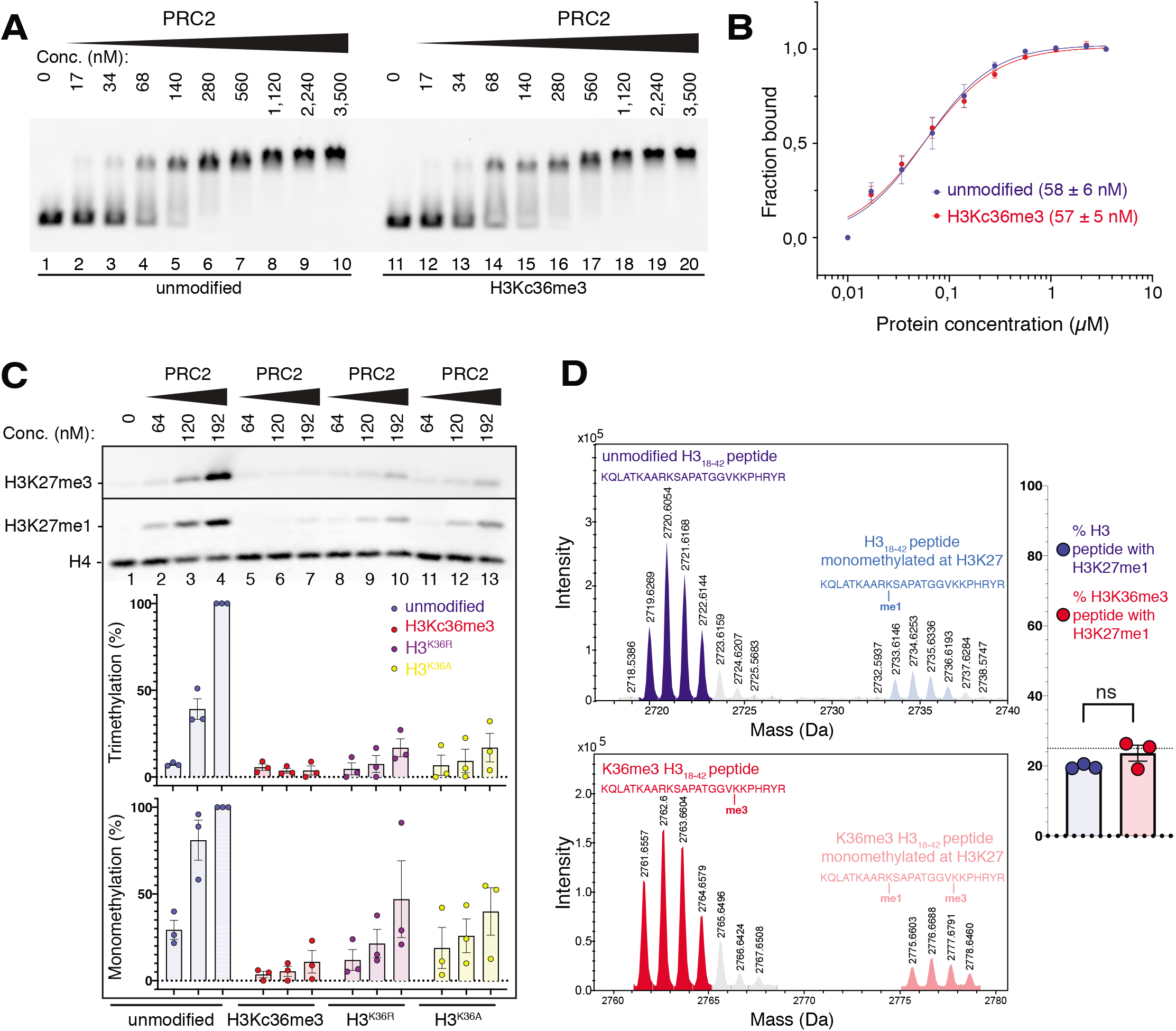
The unmodified H3K36 side chain in the EZH2_CXC_-DNA interaction interface is critical for H3K27 methylation on nucleosomes. (**A**, **B**) EMSA analysis and quantification as in **Figure 2A and B**, using PRC2 and mononucleosomes that were unmodified (lanes 1-10) or contained a trimethyllysine analog at H3K36 (H3Kc36me3, lanes 11-20). (**C**) Western Blot (WB) analysis of HMTase reactions with PRC2 as in **Figure 2C** on unmodified (lanes 1-4), H3Kc36me3 (lanes 5-7), H3^K36R^ (lanes 8-10) or H3^K36A^ (lanes 11-13) mononucleosomes (446 nM). Coomassie stained gel of reactions is shown in **Supplementary Figure 6A**. Bottom: quantification of H3K27me3 and H3K27me1 chemiluminescence signals, respectively, by densiometry analysis from three independent experiments. In each experiment, the methylation signal in lane 4 was defined as 100% and used to quantify the corresponding H3K27 methylation signals in the other lanes on the same membrane. Circles show individual data points and error bars SEM. (**D**) HMTase reactions monitoring H3K27me1 formation by PRC2 on H3_18-42_ peptides that were unmodified (top) or contained K36me3 (bottom). Left: Deconvoluted ESI-MS spectra from data shown in **Supplementary Figure 6B**. On both substrates, areas of the four colored peaks of H3K27me1-modified and unmodified substrate peptides were used for quantification of H3K27me1 formation. Right: Symbols represent percentages of peptides carrying H3K27me1 in three independent experiments, error bars show SEM; Welch’s t-test showed no significant (ns) difference between H3K27 monomethylation on the two peptide substrates.

### H3K36me3 inhibits H3K27 methylation by PHF1-PRC2

DNA-binding by the winged-helix domain of PHF1 increases the binding affinity and residence time of PHF1-PRC2 on nucleosomes about two- to three-fold, resulting in more efficient H3K27 methylation by this complex compared to PRC2 (Choi et al., 2017). Furthermore, the PHF1 tudor domain binds to H3K36me3 in the context of a nucleosome (Musselman et al., 2013). To investigate whether these interactions might modulate PRC2 inhibition by H3K36me3, we compared the HMTase activity of full-length PHF1-PRC2 (**Figure 1B**) on unmodified and H3Kc36me3 mononucleosomes. H3K27 mono- and tri-methylation by PHF1-PRC2 was strongly inhibited on H3Kc36me3 mononucleosomes (**Supplementary Figure 6C**). H3K36me3 therefore inhibits H3K27 methylation by PHF1-PRC2 even though this complex has higher binding-affinity and a prolonged residence time on nucleosomes (Choi et al., 2017). Further analyses will be needed to assess whether and how interaction of the PHF1 tudor domain with H3K36me3 might change H3K27 methylation by PHF1-PRC2 on more complex oligonucleosome substrates containing H3K36me3- and unmodified nucleosomes.

### *Drosophila* with H3^K36R^ or H3^K36A^ mutant chromatin arrest development at different stages

The observation that PRC2 is not only inhibited on H3K36me2/3-modified nucleosomes but also on H3^K36R^ and on H3^K36A^ mutant nucleosomes prompted us to investigate how H3K27 trimethylation is affected in *Drosophila* with H3^K36R^ or H3^K36A^ mutant chromatin. H3K27me3 is primarily found on canonical histone H3 (McKay et al., 2015; Pengelly et al., 2013). We used the following strategy to replace the canonical histone H3 gene copies encoded in the *HisC* gene cluster with H3^K36R^ or H3^K36A^ mutant versions. Animals that are homozygous for a deletion of the *HisC* gene cluster (i.e. *Df(2L)HisC* homozygotes) arrest development at the blastoderm stage after exhaustion of the pool of maternally-deposited histones but transgene cassettes providing 12 copies of the wild-type histone gene unit (*12xHisGU^WT^*) rescue *Df(2L)HisC* homozygotes into viable adults (Günesdogan et al., 2010; McKay et al., 2015). We therefore generated *Df(2L)HisC* homozygotes carrying *12xHisGU^H3K36R^* or *12xHisGU^H3K36A^* transgene cassettes and shall refer to these animals as *H3^K36R^* and *H3^K36A^* mutants, respectively. The *Drosophila* strains to generate *H3^K36R^* mutant animals had been described previously (McKay et al., 2015). The strain for generating *H3^K36A^* mutants was constructed in this study.

*H3^K36R^* mutant animals complete embryogenesis and their cuticle morphology is indistinguishable from wildtype (**Supplementary Figure 7**). In agreement with the results from Matera and colleagues (McKay et al., 2015), we found that these animals arrest development during the larval or pupal stages. Specifically, 81% of *H3^K36R^* mutant animals arrested development at variable time points during larval growth, 18% develop to form pupae that die prior to metamorphosis, and 1% develop into late pupae that complete metamorphosis but then arrest as pharate adults (**Supplementary Figure 7**). Like Matera and colleagues (McKay et al., 2015), we have not observed any *H3^K36R^* mutants that eclose from the pupal case, and both our studies therefore disagree with a report from the Schwartz lab who claimed that *H3^K36R^* mutants would be able to develop into adults (Dorafshan et al., 2019). When we dissected the rare *H3^K36R^* mutant pharate adults from their pupal cases and examined their epidermal structures, we found that they consistently showed homeotic transformations reminiscent of Polycomb group (PcG) mutants. These PcG mutant phenotypes included antenna-to-leg transformations and extra sex comb teeth on meso- and metathoracic legs in males (**Supplementary Figure 7**). A molecular analysis of these PcG phenotypes will be presented below.

*H3^K36A^* mutants also complete embryogenesis and the morphology of their embryonic cuticle also appeared indistinguishable from wildtype (**Supplementary Figure 7**). 96% of the *H3^K36A^* mutant animals arrest development before hatching from the eggshell and the 4% that hatch die during the first larval instar (**Supplementary Figure 7**). *H3^K36A^* mutants therefore arrest development earlier and in a narrower time window than *H3^K36R^* mutants. The molecular basis for the earlier lethality of *H3^K36R^* mutant animals remains to be determined. However, it is not unusual that mutations changing the chemical properties of a particular histone lysine residue result in phenotypes with different severity (e.g. (Copur et al., 2018))

### *Drosophila* with H3^K36R^ or H3^K36A^ mutant chromatin show diminished H3K27me3 levels at canonical PcG target genes

We next performed Western blot analyses to examine H3K36me2, H3K36me3 and H3K27me3 bulk levels in *H3^K36R^* and *H3^K36A^* mutant animals. In the case of *H3^K36R^* mutants, we used extracts from diploid imaginal disc and central nervous system (CNS) tissues dissected from third instar larvae, and in the case of *H3^K36A^* mutants we used total nuclear extracts from late-stage embryos. For the interpretation of the following experiments, it is important to keep in mind that *H3^K36R^* and *H3^K36A^* zygotic mutant animals initially also contain a pool of maternally-deposited wild-type canonical H3 molecules that, together with H3^K36R^ and H3^K36A^, become incorporated into chromatin during the pre-blastoderm cleavage cycles. Even though these wild-type H3 molecules then become diluted during every cell cycle and are eventually fully replaced by mutant H3, they are probably still present in the chromatin of late-stage embryos because of the few cell divisions that take place prior to the end of embryogenesis. In diploid tissues from larvae, replacement by mutant H3 is expected to be much more complete because of the extensive cell proliferation that occurs in these tissues during larval growth.

In *H3^K36R^* mutant larvae, H3K36me2 and H3K36me3 bulk levels were reduced more than 4-fold compared to wildtype (**Figure 4A**). The residual H3K36me2 and H3K36me3 signals (**Figure 4A**, lane 4) probably represent the methylated versions of the histone variant H3.3 that are encoded by the genes *H3.3A* and *H3.3B* that are not located in the *HisC* locus and had not been mutated in these animals. Intriguingly, *H3^K36R^* mutant animals also showed an about two-fold reduction in H3K27me3 bulk levels compared to wildtype (**Figure 4A**, compare lanes 4-6 with 1-3). The reduction of not only H3K36me2 and H3K36me3 but also of H3K27me3 bulk levels in *H3^K36R^* mutant larvae was previously also noted by Matera and colleages (Meers et al., 2017). H3K27 tri-methylation by PRC2 therefore appears to be compromised in *Drosophila* chromatin consisting of H3^K36R^ nucleosomes.

**FIGURE 4.**
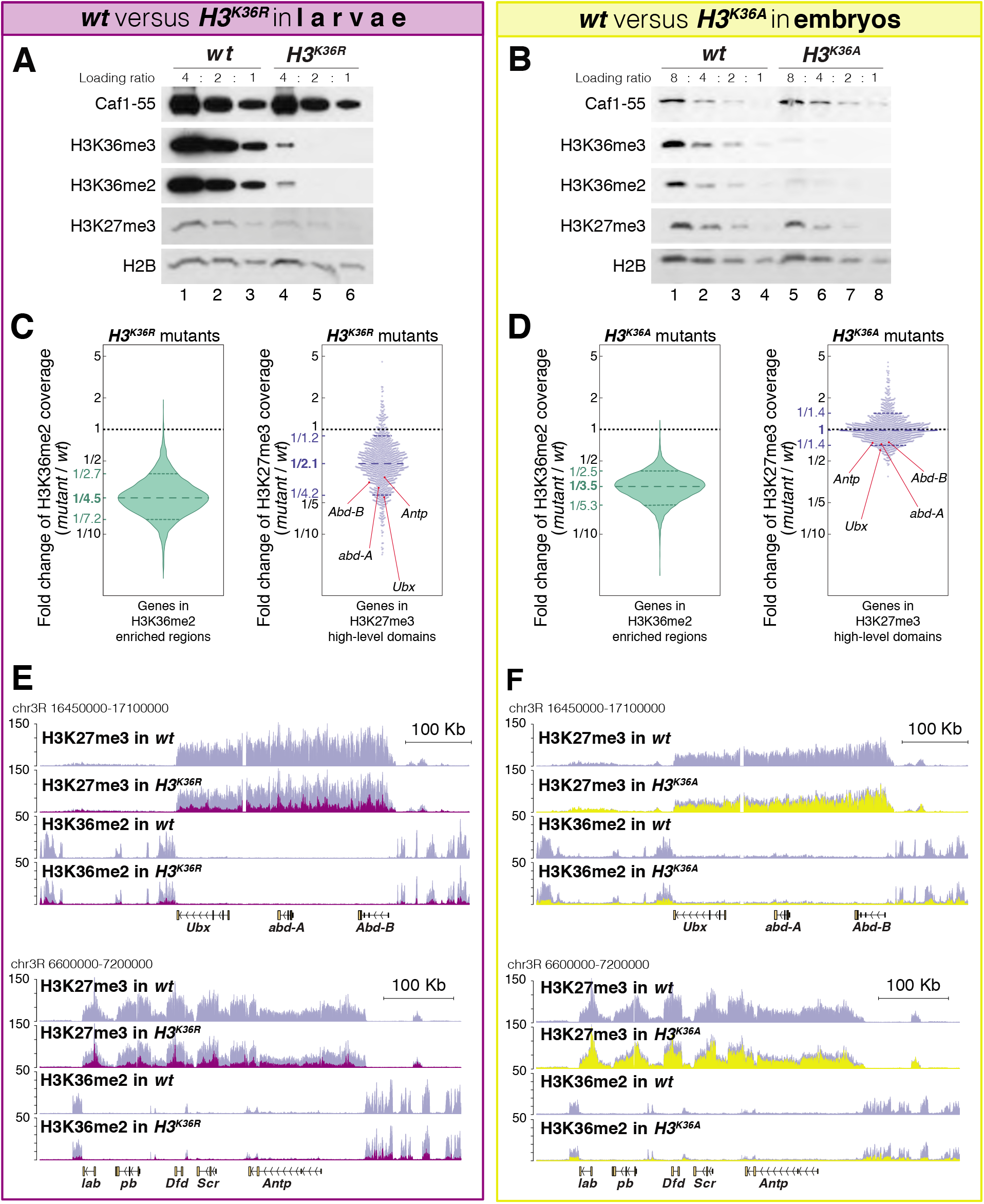
*H3^K36A^* and *H3^K36R^* mutants show reduced levels of H3K27me3. (**A**) Western blot analysis on serial dilutions (4:2:1) of total cell extracts from wing, haltere and 3^rd^ leg imaginal disc tissues dissected from *wildtype* (*wt*, lanes 1-3) and *H3^K36R^* mutant (lanes 4-6) third instar larvae. Blots were probed with antibodies against H3K36me3, H3K36me2 or H3K27me3; in each case, probing of the same membranes with antibodies against Caf1-55 and H2B served as controls for loading and western blot processing. Note the reduced levels of H3K36me3 and H3K36me2 but also of H3K27me3 in *H3^K36R^* mutants compared to *wildtype* (*wt*) (see text). See Materials and Methods for details of all genotypes. (**B**) Western blot analysis on serial dilutions (8:4:2:1) of total nuclear extracts from 21-24 hr old *wt* (lanes 1-4) and *H3^K36A^* mutant (lanes 5-8) embryos, probed with antibodies against H3K36me3, H3K36me2 or H3K27me3; and with antibodies against Caf1-55 and H2B as controls. Note that H3K36me3 and H3K36me2 levels are reduced in *H3^K36R^* mutants compared to *wt* but that H3K27me3 levels appear undiminished in the mutant (see text). (**C**) Left, violin plot showing the fold-change of H3K36me2 coverage in *H3^K36R^* mutant larvae relative to *wt* at genes that in wildtype larval CNS and imaginal disc tissues are decorated with H3K36me2 (see Materials and Methods). The dashed line marks the median reduction (4.5-fold), the dotted lines indicate indicated the interval comprising 80% of regions. Right, Bee plot showing the fold-change of H3K27me3 coverage in *H3^K36R^* mutant larvae relative to *wt* at genes that in wildype larval CNS and imaginal disc tissues are associated with high-level H3K27me3 regions (see Materials and Methods). The dashed line marks the median reduction (2.1-fold), the dotted lines indicate the interval comprising 80% of regions. Note that H3K27me3 coverage at the HOX genes *abd-A, Abd-B, Ubx* and *Antp* is between 3- and 4-fold reduced. (**D**) Analysis and representation as in (**C**) but showing fold-changes in H3K36me2 and H3K27me3 coverage in *H3^K36A^* mutant late-stage embryos relative to *wt* at genes that in wildtype embryos are decorated with H3K36me2 and H3K27me3, respectively. Note that H3K27me3 coverage at the HOX genes *abd-A, Abd-B, Ubx* and *Antp* is about 1.5-fold reduced. See also **Supplementary Figure 8**. (**E**) H3K27me3 and H3K36me2 ChIP-seq profiles in larval CNS and imaginal disc tissues from *wt* (blue) and *H3^K36R^* mutant (purple) third instar larvae; in the tracks showing the profiles in the *H3^K36R^* mutant, the *wt* profile is superimposed as reference (see **Table S2** and Materials and Methods for information about normalization). Top: genomic interval containing the *Bithorax-Complex* harbouring the HOX genes *Ubx, abd-A* and *Abd-B*; bottom: genomic interval containing the *Antennapedia-Complex* with the HOX genes *lab*, *pb*, *Dfd, Scr* and *Antp*. Note the 3- to 4-fold reduction of H3K27me3 levels across the *Bithorax* and *Antennapedia* loci in *H3^K36R^* mutants. Also note that the analyzed tissues (CNS, thoracic imaginal discs and eye-antenna discs) represent a mixed population of cells with respect to transcriptionally active and repressed states of each HOX gene. For each HOX gene there is a substantially larger proportion of cells in which the gene is decorated with H3K27me3 and repressed by Polycomb and only a small proportion of cells in which the gene is active and decorated with H3K36me2. (**F**) H3K27me3 and H3K36me2 ChIP-seq profiles at the *Bithorax* and *Antennapedia* loci as in (**E**) but from *wt* (blue) and *H3^K36A^* mutant (yellow) late-stage embryos with the *wt* profile superimposed in the tracks showing the profiles in the *H3^K36A^* mutant. H3K27me3 levels across the *Bithorax* and *Antennapedia* loci in *H3^K36A^* mutants are only about 1.5-fold reduced compared to *wt*.

In *H3^K36A^* mutant embryos, H3K36me2 and H3K36me3 bulk levels were reduced about 3- to 4-fold comprared to wildtype (**Figure 4B**, compare lanes 5-8 with 1-4). Part of the residual H3K36me2 and H3K36me3 methylation signals in *H3^K36A^* mutant embryos probably represents the methylated versions of the histone variant H3.3. However, considering that the reduction is less pronounced than in *H3^K36R^* mutant larvae, at least some of the H3K36me2 and H3K36me3 signal might also represent maternally-deposited wild-type canonical H3. H3K27me3 bulk levels appeared largely unchanged compared to wildtype (**Figure 4B**, compare lanes 5-8 with 1-4).

We next performed ChIP-seq experiments to examine how the genome-wide profiles of H3K36me2 and H3K27me3 are changed in *H3^K36R^* and *H3^K36A^* mutants. In the case of *H3^K36R^* mutants, we compared these profiles in cells from imaginal disc and CNS tissues dissected from late-stage third instar *H3^K36R^* and wildtype larvae, and in the case of *H3^K36A^* mutants we compared the profiles in late-stage *H3^K36A^* and wildtype embryos. As expected from the Western blot analyses (**Figure 4A, B**), H3K36me2 levels across the genome were strongly diminished, both in *H3^K36R^* mutant larvae and in *H3^K36A^* mutant embryos (**Figure 4C-F, Supplementary Figure 8, Table S2**). The H3K27me3 profiles confirmed that the levels of this modification were reduced in *H3^K36R^* mutant larvae (**Figure 4C**). While the average reduction was only about two-fold, H3K27me3 levels were particularly strongly diminished at canonical PRC2 target genes such as the HOX genes that in wildtype animals are decorated with high-levels of H3K27me3 (**Figure 4C, E, Supplementary Figure 8, Table S2**). Specifically, at the HOX genes *Ultrabithorax (Ubx), abdominal-A (abd-A), Abdominal-B (Abd-B*) or *Antennapedia (Antp*), H3K27me3 levels in *H3^K36R^* mutants were between 3- and 4-fold lower than in wildtype (**Figure 4C, E**). As expected from the Western blot analyses (**Figure 4B**), *H3^K36A^* mutant embryos did not show a general reduction in the H3K27me3 profile (**Figure 4D**). However, H3K27me3 levels were 1,5-fold reduced across the HOX genes (**Figure 4D, F**). In *Drosophila* with H3^K36R^ or H3^K36A^ chromatin, PRC2 therefore appears to be unable to generate high levels of H3K27me3 at Polycomb target genes.

### Polycomb repression of HOX genes is impaired in *Drosophila* with H3^K36R^ or H3^K36A^ mutant chromatin

The PcG-like phenotypes in the rare *H3^K36R^* mutant animals that survive into pharate adults and the reduction of H3K27me3 levels at HOX genes in these mutants prompted us to analyse whether and how expression of these genes is altered in *H3^K36R^* and *H3^K36A^* mutants. In a first set of experiments, we analysed HOX gene expression in embryos. Both mutants showed stochastic misexpression of *Abd-B* in single cells in late stage embryos (**Figure 5A**). *Abd-B* misexpression in *H3^K36R^* and *H3^K36A^* mutant embryos was however clearly less widespread than in *H3^K27R^* mutant embryos or in embryos lacking the PRC2 subunit Esc that are shown for comparison (**Figure 5A**). Moreover, we were unable to detect misexpression of *Antp* or *Ubx* in *H3^K36R^* or *H3^K36A^* mutant embryos.

**FIGURE 5.**
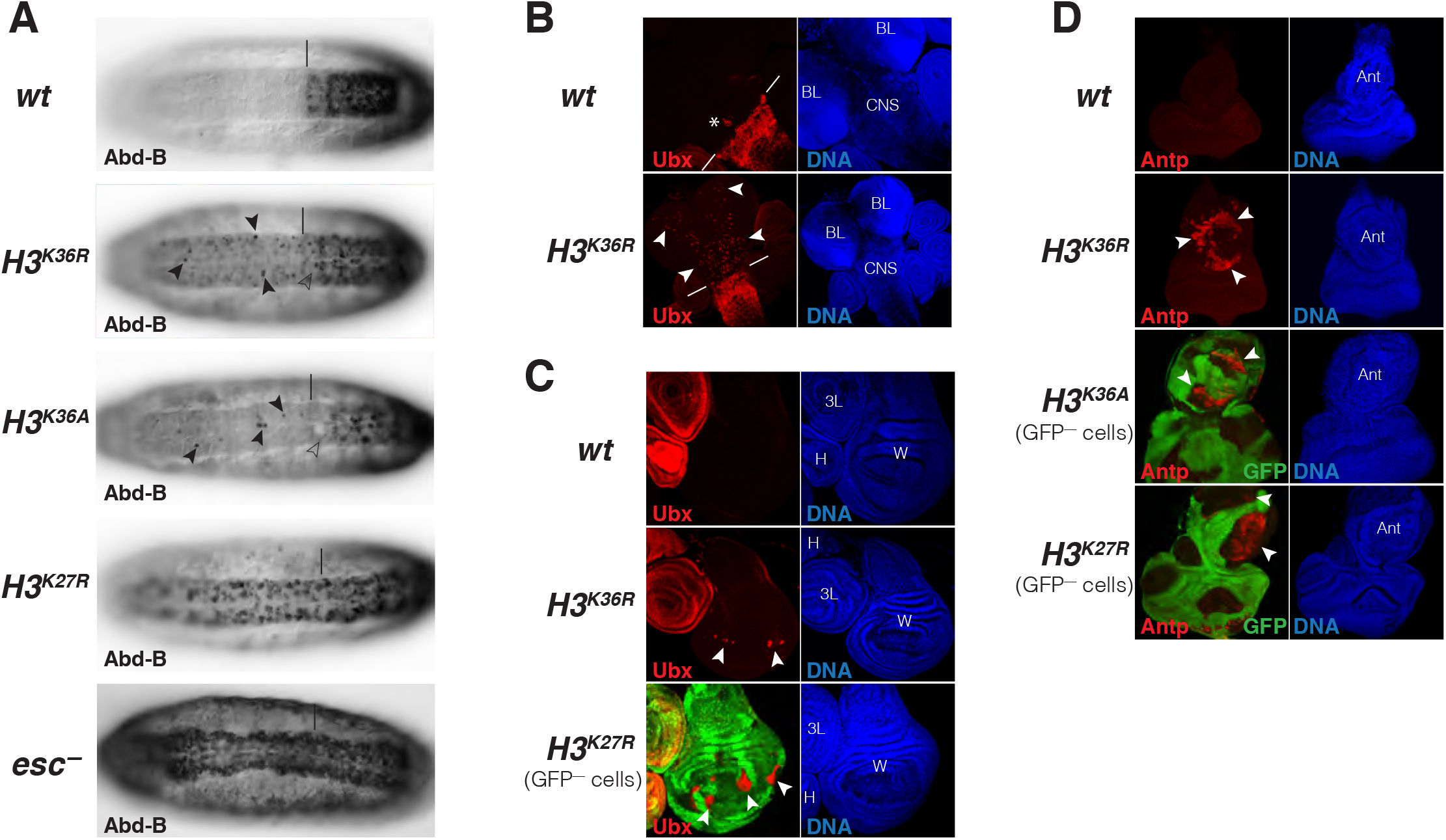
*Drosophila* with H3^K36R^ or H3^K36A^ chromatin show defective Polycomb repression at HOX genes. (**A**) Ventral views of stage 16 wildtype (*wt*), *H3^K36A^, H3^K36R^, H3^K27R^*, or *esc* (esc^−^) mutant embryos, stained with antibody against Abd-B protein; the *esc* mutant embryo lacked both maternal and zygotic expression of *esc* (see Materials and Methods for details of all genotypes). The vertical bar marks the anterior boundary of *Abd-B* expression in parasegment (ps) 10 in *wt* embryos. Note the stochastic misexpression of Abd-B protein in single cells anterior to ps10 in *H3^K36R^* and *H3^K36A^* mutant embryos (arrowheads). *H3^K27R^* and *esc* mutant embryos show widespread misexpression of Abd-B protein in the head-to-tail pattern characteristic of PcG mutants. For reasons that are not well understood, *H3^K36A^* and *H3^K36R^* mutants also show partial loss of Abd-B expression in cells in ps10 (empty arrowheads). (**B**) Larval CNS and brain lobe tissues from wildtype (*wt*) or *H3^K36R^* mutant third instar larvae, stained with antibody against Ubx protein (red) and Hoechst (DNA) to label all nuclei; location of CNS and brain lobes (BL) are indicated in the right panel. The white bars mark the anterior boundary of *Ubx* expression in ps5 in *wt* embryos, the asterisk marks the Ubx-expressing cells in the central midline of ps4 that are part of the wild-type Ubx pattern. Note the stochastic misexpression of Ubx protein in many single cells anterior to ps5 in the CNS and in the brain lobes (arrowheads). (**C**) Imaginal wing (W), haltere (H) and 3^rd^ leg (3L) discs from wildtype (*wt*) or *H3^K36R^* mutant third instar larvae and, as reference, discs from a larvae with clones of *H3^K27R^* mutant cells that are marked by the absence of GFP. In all cases discs were stained with antibody against Ubx protein (red) and Hoechst (DNA) to label all nuclei. In *wt* animals, Ubx is expressed in the halter and 3^rd^ leg disc but not in the wing disc where it is repressed by the PcG machinery. Note that in *H3^K36R^* mutants, Ubx is misexpressed in small clusters of cells in the pouch area of the wing disc (arrowheads) but remains repressed in the rest of the wing disc. Such misexpression was detected in 50% of wing discs (*n* = 28). As reference, a wing discs with *H3^K27R^* mutant clones is shown, where all cells in the clones in the wing pouch (arrowheads) show misexpression of Ubx and only mutant cells in the notum and hinge show no misexpression (empty arrowheads) (cf. (Pengelly et al., 2013)). Also note that in *H3^K36R^* mutants (*n* > 30 mutant animals analysed), Ubx expression in haltere and leg discs appears unperturbed (asterisks). (**D**) Eye-antennal imaginal discs from wildtype (*wt*) or *H3^K36R^* mutant larvae and below discs from larvae with clones of *H3^K36A^* or *H3^K27R^* mutant cells that are marked by the absence of GFP. All animals were stained with antibody against Antp protein (red) and Hoechst (DNA) to label all nuclei. Antp is not expressed in the eye-antennal disc of *wt* animals. Note that in *H3^K36R^* mutant discs, Antp is misexpressed in large clusters of cells (arrowheads) in the antenna primordium (Ant). Note that Antp is also misexpressed in *H3^K36A^* or *H3^K27R^* mutant cell clones in the antenna primordium (arrowheads) and that in these cases misexpression also only occurs in a subset of the mutant cells and not in all clones.

We next analysed HOX gene expression in imaginal discs and CNS tissues from third instar *H3^K36R^* mutant larvae. In the CNS of every single mutant individual, *Ubx* was widely misexpressed in many single cells in an apparently stochastic pattern (**Figure 5B**). 50% of the *H3^K36R^* mutant larvae also showed stochastic misexpression of *Ubx* in individual cells in the wing pouch of the wing imaginal disc (**Figure 5C**). *Ubx* misexpression in *H3^K36R^* mutant wing discs was less widespread than in *H3^K27R^* mutants that are shown for comparison (**Figure 5C**). Finally, we found that 100% of the *H3^K36R^* mutant larvae showed misexpression of *Antp* in the antenna primordium of the eye-anntennal disc (**Figure 5D**). We also observed this misexpression in clones of *H3^K36A^* homozygous cells that we had induced in *H3^K36A^* heterozygous animals (**Figure 5D**) and in *H3^K27R^* mutant clones that were induced as control (**Figure 5D**). *Drosophila* with chromatin consisting of *H3^K36R^* or *H3^K36A^* nucleosomes therefore show stochastic misexpression of multiple HOX genes. The most straightforward interpretation of this misexpression is that it is caused by defective Polycomb repression as a result of the reduced H3K27me3 levels in HOX gene chromatin.

## DISCUSSION

Understanding how PRC2 binds chromatin and how it is regulated is essential for understanding how the complex marks genes for Polycomb repression to maintain cell fate decisions. The work in this study leads to the following main conclusions. First, the structure of nucleosome-bound PHF1-PRC2 allowed to visualize how interaction of the catalytic lobe of the complex with the substrate nucleosome threads the histone H3 N-terminus into the active site of EZH2 through a relay of contacts. Second, structure-guided mutational analyses showed that DNA-binding by the EZH2_CXC_ domain is critical for productive PRC2-nuclesome interactions. Third, unmodified H3K36 is accommodated in a key position in the EZH2_CXC_-DNA interface and while H3K36 provides the correct fit, the methylated forms H3K36me2/3, or mutated H3K36R or H3K36A do not seem to fit because they strongly diminish H3K27 methylation. Fourth, H3K36 is also critical for normal H3K27 methylation *in vivo* because *Drosophila* with H3^K36R^ or H3^K36A^ mutant chromatin show reduced levels of H3K27me3 and fail to fully maintain Polycomb repression at HOX target genes. In the following, we shall discuss key aspects of these new findings in the context of our previous knowledge of PRC2 regulation and function.

### Different forms of PRC2 use the same molecular interactions for binding the H3 N-terminus on substrate nucleosomes

Unlike many other histone-modifying enzymes (e.g. (McGinty et al., 2014; Worden et al., 2019)), PRC2 does not recognize the nucleosome by docking on its acidic patch (Luger et al., 1997) to engage with the histone substrate. Instead, the complex interacts with chromatin by binding to the DNA gyres on the nucleosome ((Poepsel et al., 2018), this study). Prevous studies that had measured the binding affinity and residence time of PRC2 on nucleosomes and free DNA had found that DNA-binding makes the largest make contribution to the chromatin-binding affinity of PRC2 (Choi et al., 2017; Wang et al., 2017). The mutational analyses here establish that interaction of highly conserved residues in the EZH2_CXC_ domain with the DNA on the substrate nucleosome is critical for H3K27 methylation (**Figure 2C**). Moreover, this interaction sets the register for a network of interactions of the H3 N-terminus with the EZH2 surface that permits H3K27 to reach into the active site (**Figure 1 D, F**). Consistent with our findings here, an independent recent study of a cryo-EM structure of PRC2 with co-factors JARID2 and AEBP2 bound to a mononucleosome with monoubiquitylated H2A (Kasinath et al., 2020) identified very similar interactions of EZH2 with the nucleosomal DNA and the H3 N-terminus. Different forms of PRC2 that contain different accessory proteins and dock in different ways on chromatin therefore contact the substrate H3 N-terminus in the nuclesosome through similar interactions.

### The position of H3K36 in the EZH2_CXC_-nucleosome interface enables allosteric regulation by H3K36 methylation

Important novel insight from our structure came from the observation that unmodified H3K36 is located in a critical position in the EZH2_CXC_-DNA interface. Unmodified H3K36 has the right fit for interaction of the H3 N-terminus with the EZH2 surface and placement of H3K27 in the active site. The inhibition of H3K27 mono-, di- and tri-methylation on nucleosomes carrying H3K36me2 or -me3 (Schmitges et al., 2011; Yuan et al., 2011) or on H3^K36R^ or H3^K36A^ nucleosomes (**Figure 3C**) suggests that these alterations of the H3K36 side chain impair the interaction of H3K27 with the active site of EZH2. On isolated H3 N-terminal peptides, H3K36me3 did not inhibit the formation of H3K27me1 (**Figure 3D**), consistent with earlier findings that on peptide substrates H3K36me3 only has a minor effect on the kcat of H3K27 methylation (Schmitges et al., 2011) (Jani et al., 2019). Also, H3K36me3 does not diminish the affinity of PRC2 for binding to mononucleosomes (**Figure 3A, B**) and does not reduce the residence time of PRC2 on nucleosome arrays (Guidotti et al., 2019). Taken together, a possible scenario would therefore be that within the time frame of the PRC2 nucleosome binding and reaction cycle, docking of the H3K36 side chain in the EZH2_CXC_-DNA interface is critical for rapid alignment of the H3 N-terminus on the EZH2 surface into a catalytically competent state. According to this view, H3K36me2/3 does not locally disrupt nucleosome binding but allosterically inhibits H3K27 from interacting with the EZH2 active site.

### H3K27 methylation and Polycomb repression are defective in Drosophila with H3^K36R^ or H3^K36A^ chromatin

The finding that PRC2 is inhibited on H3^K36R^ and H3^K36A^ nucleosomes *in vitro* had prompted us to use a genetic histone replacement strategy in *Drosophila* (Günesdogan et al., 2010) (McKay et al., 2015) to assess PRC2 inhibition on H3^K36R^ or H3^K36A^ chromatin *in vivo*. Previous studies had found that *Drosophila H3^K36R^* mutants are able to develop into the pupal stages and, consistent with this late develpomental arrest, whole third instar larvae were found to show only relatively minor changes in their transriptome compared to wildtype animals (McKay et al., 2015; Meers et al., 2017). Here, we found that a few rare *H3^K36R^* mutant animals even survive into pharate adults and that these show remarkably little morphological defects apart from homeotic transformations characteristic of Polycomb mutants (**Figure S7**). We show that these phenotypes are caused by defective Polycomb repression of multiple HOX genes (**Figure 5**) and that they are linked to reduced levels of H3K27me3 at these genes (**Figure 4**). A simple straightforward explanation for these phenotypes in *H3^K36R^* or *H3^K36A^* mutant animals is that PRC2 is unable to effectively deposit high levels of H3K27me3 on the H3^K36R^ or H3^K36A^ nucleosomes, respectively, in their chromatin. Accordingly, H3K27me3 levels at HOX genes are below the threshold needed to reliably maintain Polycomb repression and consequently, HOX genes become stochastically misexpressed in a fraction of cells. Finally, we note that in *H3^K36R^* mutant larvae, the experimental setting where we have been able to generate the most complete replacement of H3 by H3^K36R^, H3K27me3 levels at HOX genes were only about 3- to 4-fold reduced compared to wildtype (**Figure 4C**). However, as shown in **Figure 3C**, on nucleosomes *in vitro*, H3K36me3 inhibited PRC2 more effectively than H3^K36R^ or H3^K36A^. It therefore seems likely that in contrast to the *H3^K36R^* and *H3^K36A^* mutants that we have used as proxy, H3K36me2 and H3K36me3 *in vivo* also inhibit PRC2 more effectively from depositing H3K27me3 on H3K36me2- or H3K36me3-modified nucleosomes in transcriptionally active chromatin.

### Concluding remark

The structural, biochemical and genetic work reported in this study shows that it is the exquisite geometry formed by a relay of interactions between the PRC2 enzyme, nucleosomal DNA and the H3 N-terminus that enable the histone methylation marks H3K36me2 and H3K36me3 in transcriptionally active chromatin to allosterically prevent PRC2 from depositing the repressive histone methylation mark H3K27me3 at transcribed genes.

## MATERIALS AND METHODS

### Protein expression and purification

Human PHF1-PRC2 wild-type (wt) complex was expressed and purified as previously described (Choi et al., 2017). In brief, an optimized ratio of the baculoviruses for the different PHF1-PRC2 subunits was used to infect HiFive cells (Invitrogen). Cell were lysed using a glass Dounce homogenizer and the complex was purified using affinity chromatography (Ni-NTA and Strep-tag), followed by simultaneous TEV mediated protease tag cleavage and Lambda Phosphatase treatment (obtained from the MPI of Biochemistry Protein Core facility) and a final size exclusion chromatography (SEC) step in a buffer containing 25 mM Hepes, pH 7.8, 150 mM NaCl, 10% glycerol, 2 mM DTT.

PRC2^CXC>A^, PRC2^EED>A^ and PRC2^CXC>A/EED>A^ mutants were generated by PCR with primers containing the desired mutations, subsequent ligation and transformation. Expression and purification were performed as above.

*Xenopus laevis* (*X.l*.) and *Drosophila melanogaster* (*D.m*.) histones were expressed and purified from inclusion bodies as described in (Luger et al., 1999). To mimic the inhibitory mark H3K36me3 or the allosteric activating mark H3K27me3, the cysteine side chain of a mutated *D.m*. histone H3^C110A K36C^ or *X.l*. histone H3^C110A K27C^ was alkylated with (2-bromoethyl) trimethylammonium bromide (Sigma Aldrich) as described previously (Simon et al., 2007). Nucleosomes containing these modifications are abbreviated with e.g. H3Kc36me3.

For histone octamers, equimolar amounts of histones H2A, H2B, H4 and H3 (wt, H3^K36A^, H3^K36R^, H3Kc27me3 or H3Kc36me3) were mixed and assembled into octamers in high salt buffer containing 10 mM Tris-HCL pH 7.5, 2 M NaCl, 1 mM EDTA, 5 mM β-mercaptoethanol. Subsequent SEC was performed to separate octamers from H3/H4 tetramers or H2A/H2B dimers (Luger et al., 1999).

#### Reconstitution of <u>nucleosomes</u>

For *X.l*. and *D.m* <u>mononuclesomes</u> used in biochemical assays, 6-carboxyfluorescein (6-FAM)-labeled 215 bp 601 DNA (Lowary and Widom, 1998) was PCR amplified from the p601 plasmid, purified on a MonoQ column (GE Healthcare), precipitated with ethanol and dissolved in the same high salt buffer used for octamers. Optimized ratios of octamer to DNA (usually ranging between 0.8-1.3 : 1) were mixed and nucleosomes were reconstituted by gradient and stepwise dialysis against low salt buffers to a final buffer containing 25 mM Hepes, pH 7.8, 60 mM NaCl, 2 mM DTT.

*X.l*. <u>asymmetrical dinucleosomes</u> for cryo-EM studies containing one unmodified substrate nucleosome and one H3K27me3-modified (allosteric) nucleosome connected with a 35 bp linker DNA were reconstituted using the protocol described in (Poepsel et al., 2018). In brief, substrate nucleosomes and allosteric nucleosomes were separately assembled on the respective *DraIII* digested nucleosomal DNA. The latter was generated by PCR with primers introducing the desired linker and *DraIII* recognition sites and purified as described above. The assembled nucleosomes were purified on a preparative native gel system (Biorad 491 prep cell). After ligation using T4 ligase (Thermo Fisher Scientific) the resulting dinucleosomes were purified from aberrant or non-ligated mononucleosomes by a second preparative native gel system (Biorad 491 prep cell). In contrast to (Poepsel et al., 2018), the dinucleosome DNA used in this study contained an additional 30 bp overhang on the substrate nucleosome, thus resulting in the following DNA sequence:

5’–601 binding (allosteric nucleosome) – agcgatctCACCCCGTGatgctcgatactgtcata – 601 binding (substrate nucleosome) – atgcatgcatatcattcgatctgagctcca −3’ (after DraIII digestion, assembly of substrate/allosteric nucleosome and ligation to dinucleosomes).

*X.l*. <u>symmetrical unmodified dinucleosomes</u> used for the HMTase assays with the PRC2^CXC^ mutants were obtained by reconstituting octamers with a 377 bp DNA containing two 601 sequences connected by a 35 bp linker DNA. A vector containing the 377 bp sequence was ordered from Invitrogen GeneArt and was used for PCR resulting in:

5’–atatctcgggcttatgtgatggac – 601 binding (substrate nucleosome 1) – agcgatctcaacgagtgatgctcgatactgtcata – 601 binding (substrate nucleosome 2) – gtattgaacagcgactcgggatat–3’.

The PCR products were purified as described above. Optimized ratios of octamer : DNA (usually ranging between 1.8-2.3 : 1) were mixed and nucleosomes were reconstituted by gradient and stepwise dialysis against low salt buffers to a final buffer containing 25 mM Hepes, pH 7.8, 60 mM NaCl, 2 mM DTT.

### Cryo-EM Data acquisition

Complexes of PHF1-PRC2 and asymmetrically modified 35 bp dinucleosomes were assembled and grids were prepared as described previously, with the difference of using 0.005% NP40 instead of 0.01% (Poepsel et al., 2018). Cryo-EM data were collected on an FEI Titan Krios microscope operated at 300 kV and equipped with a post-column GIF and a K2 Summit direct detector (Gatan) operated in counting mode. A total of 3467 movies were collected at a nominal magnification of 81,000x (1.746 Å/pixel) at the specimen level using a total exposure of 53 e^-^ / Å^2^ distributed over 60 frames and a target defocus range from 1.5–3 μm. Data acquisition was carried out with SerialEM.

### Cryo-EM Data processing

Movies were aligned and corrected for beam-induced motion as well as dose compensated using MotionCor2 (Zheng et al., 2017). CTF estimation of the summed micrographs was performed with Gctf (Zhang, 2016) and particles were picked in Gautomatch (http://www.mrc-lmb.cam.ac.uk/kzhang/ K. Zhang, MRC LMB, Cambridge, UK) using templates created from the AEBP2-PRC2-dinucleosome cryo-EM structure (EMD-7306, (Poepsel et al., 2018). All subsequent image processing steps were performed in Relion 3.0 (Zivanov et al., 2018) as shown in Fig. S2. A total of 1,028,229 candidate particles were subjected to two rounds of initial 3D classification against a reference map (AEBP2-PRC2-dinucleosome low-pass filtered to 60 Å) and the Bayesian fudge factor (T value) set to 8. 330,482 remaining particles were subjected to two more rounds of 3D classification, this time using the best 3D model from the previous run as reference. Finally, the two best 3D models were 3D refined and further classified into 10 classes without translational and rotational sampling, using a T value of 4. From this run, the best 3D classes with the highest nominal overall resolution and rotational and translational accuracies were subjected to iterative rounds of 3D refinement, this time applying a soft mask for solvent flattening, per particle CTF refinement and Bayesian polishing. The highest nominal resolution was only achieved by combining several models from the previous 3D run, likely due to missing particle views in one or the other individual model. The final map after postprocessing had an overall nominal resolution of 5.2 Å, as determined from the gold-standard FSC criterion of 0.143 (Rosenthal and Henderson, 2003) (Fig. S1D). The density (Overall PHF1-PRC2:di-Nuc) with fitted models is shown in Fig.1A and in Fig. S1E using UCSF ChimeraX (Goddard et al., 2018). Local resolution estimation was performed in Relion 3.0 and is shown in Fig. S1B. The spherical angular distribution of all particles in the final model is shown in Fig. S1C.

To further improve the resolution and map details of the region around the H3 N-terminus, particle subtraction and focused 3D refinement was applied (Bai et al., 2015; Ilca et al., 2015; Zhou et al., 2015). Using a mask generated with UCSF Chimera (Pettersen et al., 2004) and Relion 3.0 the signal of the allosteric nucleosome as well as parts of PRC2 (EED and EZH2_allo_) was subtracted from all particle images. These signal subtracted particles were then subjected to focused 3D refinement using a soft mask around the substrate nucleosome and EZH2_sub_. This yielded a 4.4 Å map (EZH2_sub_-Nuc_sub_) (Fig. S3B). Local resolution estimation is shown in Fig.S3A. For model building and depiction, the final density was further sharpened (applied b – factor: - 66) using the Multisharpen function in Coot (Emsley et al., 2010) (e.g. in Figs. 1E, S3D, E and F).

To confirm the side chain information visible in the Coot sharpened map, Phenix Resolve density modification was run on the two half maps generated from the 3D refinement of the EZH2_sub_-Nuc_sub_ map (Terwilliger et al., 2019). The resolution of the map according to Phenix cryo EM density modification output improved to 4 Å and the resulting map was used as an additional guideline for model building as well as for depiction (in Figs. S4_A-D.).

### Cryo-EM data fitting, modeling and refinement

Available crystal structures were fitted into the final maps using rigid-body fitting in UCSF Chimera and all manual remodeling, morphing and building was performed in Coot. For PRC2, the crystal structure of the catalytic lobe of human PRC2 (PDB: 5HYN (Justin et al., 2016)) was used. Since the SBD helix and the SANT1 helix bundle of the crystal structure was not accommodated well by the corresponding EM density, this region was fitted separately. A model of a dinucleosome with linker DNA (Supplementary dataset 1 in (Poepsel et al., 2018), including crystal structures of nucleosomes, PDB 3LZ1, also PDB 1AOI, also PDB 6T9L) was fitted.

The above described overall model was then used as a starting model for fitting and building EZH2_sub_-Nuc_sub_ into the focused map. Where possible, missing parts in the model were built de-novo, i.e. the H3 N-terminal tail (residues 30-37) between the catalytic site of PRC2 and the substrate histone. Available information from crystal structures was used as a guide (PRC2 with H3 peptide bound: PDB: 5HYN (Justin et al., 2016), and high resolution crystal structures of nucleosomes (PDB 1AOI and PDB 6T9L) (Luger et al., 1997). Parts of EZH2_sub_-Nuc_sub_ model were then fitted using the morph fit routine in Coot or manually (Casañal et al., 2020). Secondary structure restraints for real-space refinement were generated automatically with phenix.secondary_structure_restraints (Sobolev et al., 2015) and manually curated. Hydrogens were added and the model was real-space refined with a resolution-cutoff of 4.4 Å with Phenix (Afonine et al., 2018) (phenix-1.18rc1-3777), using reference structures (PDB 6T9L and PDB 1AOI for nucleosome and one copy of the human PRC2 crystal structure generated from PDB 5HYN), applying strict secondary structure and Ramachandran restraints.

Our final model includes the modelled side chains of the fitted crystal/cryo-EM structures. This is in our opinion supported by the data as the substrate nucleosome protein core is resolved to app. 4 Å (Fig. S3A) and the map in these regions shows clear bulky side chain information (Fig S3D). The EZH2 density is of worse quality however even at lower resolution side chains likely contribute to the signal in the particle images and thereby an overall good model to map fit (in our case given by the high CC values as well as FSC_modelvsmap_) is arguably only ensured in the presence of side chains. However we caution readers against in interpreting our model at side chain resolution in poorly resolved regions.

Structures were visualized with UCSF ChimeraX (Goddard et al., 2018) and PyMOL2 (https://pymol.org/2/).

### Electrophoretic mobility shift assay (EMSA)

EMSAs on a 1.2% agarose gel in 0.4x TBE Buffer with 45 nM 6-FAM - labeled mononucleosomes (unmodified wt *X.l*. for bandshifts with the PRC2^CXC^ mutants, unmodified wt *D.m*. and *D.m* H3Kc36me3 trimethyllysine analog containing nucleosomes) and increasing PRC2 concentrations (concentrations indicated in the figures above the gels) were performed in triplicates as described in (Choi et al., 2017). A Typhoon FLA 9500 scanner and the Fiji software was used for densitometric analysis of the 6-FAM signal (Schindelin et al., 2012). Background correction and calculation of the fractions of bound nucleosomes was performed with R using tidyverse (https://www.r-project.org/). In detail: two parts were boxed out in each lane: 1. unbound nucleosomes (‘unbound’ box) and 2. shifted nucleosomes (‘bound’, everything above ‘unbound’). The boxed-out signals were integrated and background corrected by subtracting the respective control (‘bound’ background of lane 1 for ‘bound’ boxes and ‘unbound’ background of lane 10 for ‘unbound’ boxes). To calculate the fraction of bound vs. unbound nucleosomes, the value for ‘bound’ nucleosome in each lane was divided by the total signal (sum of bound and unbound) of the same lane. Hill function fitting and illustration of the plot were subsequently performed with Prism 8 (GraphPad).

### Histonemethyltransferase (HMTase) assay

For all HMTase assays, 446 nM of mononucleosomes or 223 nM of dinucleosomes were incubated with indicated amounts of the different PRC2 complexes, in a reaction buffer containing 20 mM HEPES pH 7.8, 50 mM NaCl, 2.5 mM MgCl_2_, 5% glycerol, 0.25 mM EDTA, 0.5 mM DTT and 80 μM S-adenosylmethionine (SAM). Reactions were allowed to proceed for 90 min at RT before quenching by the addition of 1x (final concentration) SDS loading buffer and heat inactivation at 95 °C for 5 min. Proteins were separated by electrophoresis on a 16% (w/v) SDS gel, transferred to a nitrocellulose membrane and probed with antibodies against H3K27me3 (Millipore, 07-449), H3K27me1 (Millipore, 07-448) and H4 (Abcam, ab10158). For quantification, HMTase reactions and the corresponding western blots on *D.m*. unmodified, H3Kc36me3, H3^K36A/R^ mononucleosomes were performed in triplicates and subjected to densitometric analysis (Chemiluminescence signal, ImageQuant LAS 4000). The integrated densitometric signal (band) in each lane was background corrected against the control lane (lane 1, no PRC2 in the reaction) and normalized with respect to the lane containing the highest amount (i.e. 100%) of PRC2 on unmodified nucleosomes (lane 4). The relative amounts of trimethylation/monomethylation for all other lanes were calculated with respect to lane 4. Graphical representations were made with Prism 8 (GraphPad).

### Mass Spectrometry (MS)

500 nM of PRC2 were incubated with 2 μM of either unmodified or H3_18-42_ peptide containing the K36me3 modification in HMTase reaction buffer (described above) and methyltransferase activity was allowed to proceed over night at RT. Reactions were then quenched with 1% trifluoroacetic acid (TFA). Home-made stage tips with poly(styrenedivinylbenzene) copolymer (SDB-XC) were used to remove PRC2 from the reactions (Rappsilber et al., 2007). First, stage tips were washed with methanol, followed by a second wash with buffer B (0.1% (v/v) formic acid, 80% (v/v) acetonitrile). The SDB-XC material was then equilibrated with buffer A (0.1% (v/v) formic acid) and 40 μl of sample was applied and washed several times. Finally, samples were eluted using buffer B and introduced into the Bruker maXis II ETD mass spectrometer by flow injection of 20 μl sample using an Agilent HPLC at a flow rate of 250 μl/min and 0.05% TFA in 70% acetonitril:H2O as solvent for ESI-MS time-of-flight analysis.

Peptides were ionized at a capillary voltage of 4500 V and an end plate offset of 500 V.

Full scan MS spectra (200-1600 m/z) were acquired at a spectra rate of 1 Hz and a collision cell energy of 15 eV.

Raw data files were processed using Bruker Compass DataAnalysis. The m/z spectra were deconvoluted (maximum entropy method) with an instrument resolving power of 10,000 and the resulting neutral spectra peaks were integrated. For quantification, the experiment was performed in triplicates. The sum of the monomethylation peak areas was divided by the sum of the first 4 peaks of the input peptide together with the sum of the monomethylation peak areas. Illustration of the quantification was subsequently performed with Prism 8 (GraphPad). A Welch’s t-test was calculated to show the nonsignificant difference between the activity of PRC2 on unmodified or H3K36me3 peptide.

### Construction of histone transgenes to generate *H3^K36A^* and *H3^K36R^* strains

Site directed mutagenesis on *pENTR221-HisGU. WT, pENTRL4R1-HisGU. WT* and *pENTRR2L3-HisGU.WT* (Günesdogan et al., 2010) was used to mutate histone H3K36 to alanine or arginine. The final constructs *pfC31-attB-3xHisGU.H3K36A* and *pfC31-attB-3xHisGU.H3K36R* were generated by Gateway LR recombination of above vectors and integrated at attP sites VK33 (BDSC 9750) and 86Fb (BDSC 130437). The full genotypes of animals used in the study is described below.

### *Drosophila* strains and genotypes

The following strains were used in this study:

*Oregon-R*
*w; Df(2L)His^C^ FRT40A/Df(2L)His^C^ FRT40A; 12xHisGU^wt^/ 12xHisGU^wt^* (McKay et al., 2015)
*w; Df(2L)His^C^ FRT40A/CyO ubi-GFP; 12xHisGU^H3K36R^/TM6B* (McKay et al., 2015)
*w; Df(2L)His^C^ FRT40A/ CyO twi:Gal4 UAS:GFP; 3xHisGU^H3K36A^(VK33) 3xHisGU^H3K36A^(86Fb)/ 3xHisGU^H3K36A^(VK33) 3xHisGU^H3K36A^(86Fb)* (generated in this study)
*w; Df(2L)His^C^ FRT40A/CyO ubi:GFP; 3xHisGU^H3K27R^(68E) 3xHisGU^H3K27R^ (86Fb)/ 3xHisGU^H3K27R^ (68E) 3xHisGU^H3K27R^ (86Fb*) (Pengelly et al., 2013)
*w hs-flp; w; hs-nGFP FRT40A/hs-nGFP FRT40; 3xHisGU^H3K27R^(68E) 3xHisGU^H3K27R^(86Fb)/3xHisGU^H3K27R^(68E) 3xHisGU^H3K27R^ (86Fb)* (Pengelly et al., 2013)
*w hs-flp; M(2)25A ubi-GFP FRT40A/CyO*
*yw; esc^6^ b pr / CyO, P[esc^+^]*
*In(2LR) Gla / CyO, esc^2^*

The following genotypes were used for the experiments shown in:

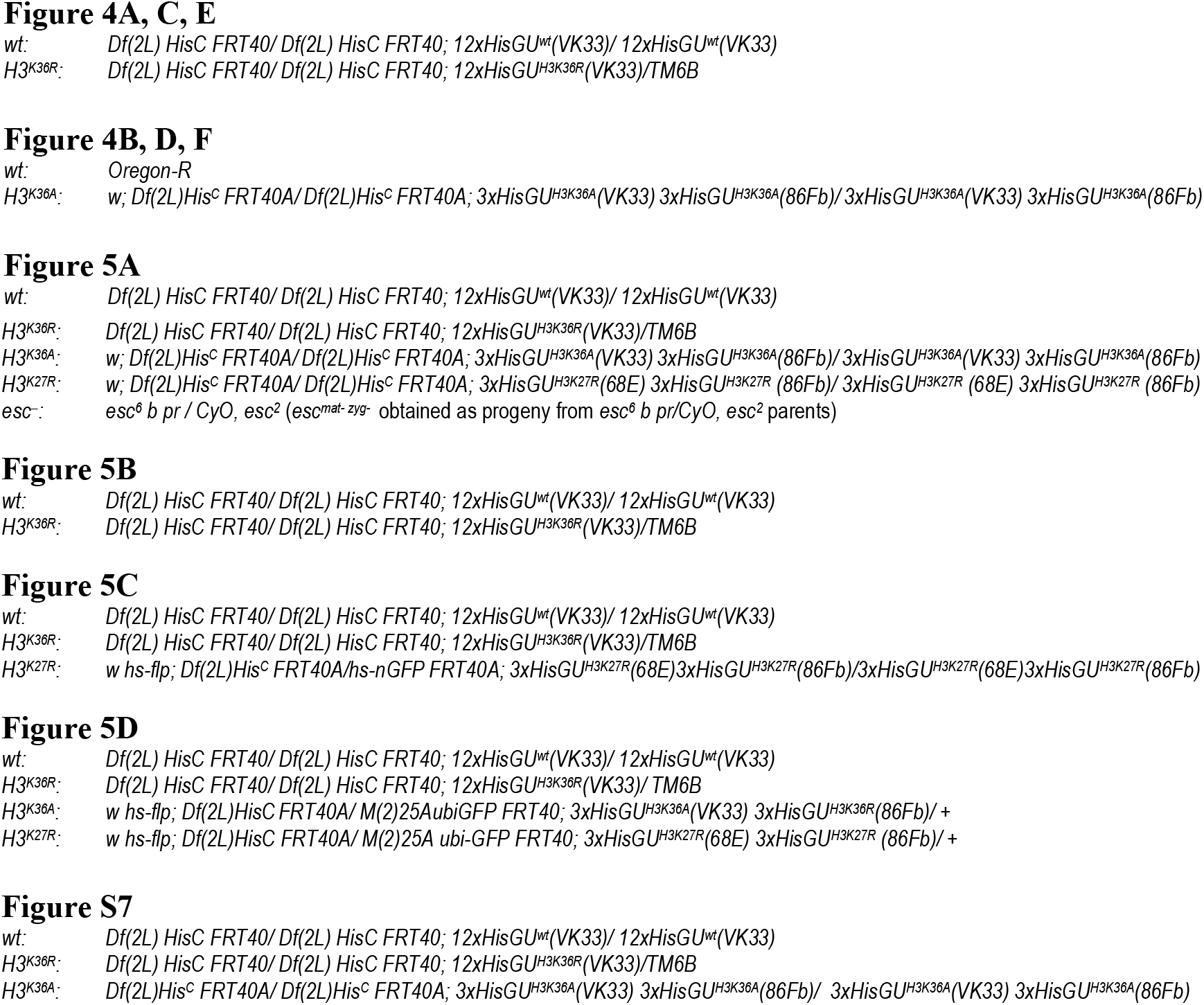

### Immunohistochemistry and immunofluorescence stainings

Embryos of the appropriate genotypes listed above were identified by the lack of GFP marked balancer chromosomes, fixed and stained with Abd-B antibody, following standard protocols. Imaginal discs from third instar larvae were stained with Antp and Cy3-labeled secondary antibodies following standard protocols. For clonal analysis (**Fig. 3D**), clones were induced 96 hrs before analyses by heat-shocked induced expression of Flp recombinase in the genotypes listed above.

### ChIP-seq analysis in *Drosophila* embryos and in larval tissues

#### Embryo collection, chromatin preparation and ChIP

_21-24 hr old *wt, H3^K36A^* embryos (see above for details of genotypes) were dechorionated, quick-frozen in liquid N2 and stored at −80°C. 5 μL of thawed embryos were homogenized in 5 mL of fixing solution (60 mM KCl, 15 mM NaCl, 4 mM MgCl_2_, 15 mM Hepes pH 7.6, 0.5% Triton X-100, 0.5 mM DTT, protease inhibitors, 0.9% Formaldehyde) at r.t.. The homogenate was filtered through a strainer (Greiner Bio-One, EASYstrainer™ 100 μm, #542 000) and incubated for 10 min with frequent gentle shaking. Cross-linking was stopped by the addition of 450 μL of 2.5 M Glycine. Fixed nuclei were washed with 1 mL of buffer A1 (60 mM KCl, 15 mM NaCl, 4 mM MgCl_2_, 15 mM Hepes pH 7.6, 0.5% Triton X-100, 0.5 mM DTT, protease inhibitors), washed with 1 mL of pre-lysis buffer (140 mM NaCl, 15 mM Hepes pH 7.6, 1 mM EDTA, 0.5 mM EGTA, 1% Triton X-100, 0.5 mM DTT, 0.1% Na Deoxycholate, protease inhibitors), resuspended in 1 mL of lysis buffer (140 mM NaCl, 15 mM Hepes pH 7.6, 1 mM EDTA, 0.5 mM EGTA, 1% Triton X-100, 0.5 mM DTT, 0.1% Na Deoxycholate, protease inhibitors, 0.1% SDS, 0.5% N-laurylsarcosine), incubated at least 10 min at 4°C with shaking, and transferred into milliTUBES 1 mL AFA Fiber (100) (Covaris, #520130) for sonication. Sonication was performed in a Covaris S220 AFA instrument using the following setup: 140W (peak incident power) / 5% (duty cycle) / 200 (cycle per burst) / 15 min. Insoluble material was removed by centrifugation in an Eppendorf centrifuge at 14000 rpm (10 min at 4°C). Input chromatin was quantified by measuring DNA concentration after decrosslinking using Qubit (Thermo Scientific) and 250 ng of chromatin were used for each ChIP experiment. 250 ng of an independently prepared batch of *D. pseudoobscura* chromatin were spiked-in in each ChIP experiment for subsequent normalization of the ChIP-seq datasets. The rest of the ChIP protocol was performed as described (Bonnet et al., 2019). For each condition, the ChIP experiment was performed in duplicates from two biologically independent chromatins. ChIP on hand-dissected CNS and imaginal disc tissues from 3^rd^ instar *wt* or *H3^K36R^* homozygous larvae (see above for details on genotypes) was performed as described (Laprell et al., 2017) with the difference *D. pseudoobscura* chromatin was spiked in at a 1:1 ratio of dm / dp chromatin.

#### Library preparation and sequencing

Library preparation for sequencing was performed with TruSeq kits from Illumina. Illumina systems (NextSeq 500) were used for paired-end DNA sequencing. All reads were aligned using STAR (Dobin et al., 2013) to the *D. melanogaster* dm6 genome assembly (Santos et al., 2015) and to the *D. pseudoobscura* dp3 genome assembly (Nov. 2004, FlyBase Release 1.03). Only sequences that mapped uniquely to the genome with a maximum of two mismatches were considered for further analyses.

#### Identification of H3K36me2 and H3K27me3 enriched regions

The Bioconductor STAN-package (Zacher et al., 2017) was used to define the location of H3K36me2-enriched regions. The seven chromosome arms (X, 2L, 2R, 3L, 3R, 4 and Y) defined in the dm6 genome assembly were segmented in 200 bp bins. STAN annotated each of these bins into 1 of 3 ‘genomic states’ based on the number of H3K36me2 ChIP-seq reads and the number of input reads overlapping with each bin, in 21-24 hr wild-type embryos. These 3 ‘genomic states’ corresponded to: ‘H3K36me2 enriched’ regions; ‘low or no H3K36me2’ regions and ‘no input’ regions. The Poisson Lognormal distribution was selected and fitting of hidden Markov models was performed with a maximum number of 100 iterations. Stretches of consecutive bins annotated as ‘H3K36me2 enriched’ regions were sometimes separated by a few bins showing another type of annotation (i.e. ‘no input’). To define a relevant set of H3K36me2 enriched regions, we considered that if stretches of consecutive bins annotated as ‘H3K36me2 enriched’ regions are not separated by more than 7 Kb, they can be fused. High-level H3K27me3 domains previously defined using the same Bioconductor STAN-package in Bonnet et al (Bonnet et al., 2019) were used in this study.

#### Normalization and visualisation of H3K27me3 and H3K36me2 ChIP-Seq datasets

The proportion of *D. pseudoobscura* reads as compared to *D. melanogaster* reads in input and in samples was used to normalize the H3K36me2 and H3K27me3 ChIP-seq datasets from *H3^K36A^* and *H3^K36R^* mutants to the corresponding wild-type H3K36me2 and H3K27me3 ChIP-seq datasets respectively (see Table S2). Chip-seq tracks shown in Fig. 4 show the average of the two biological replicates that were performed for each condition. Y-axes of ChIP-seq tracks correspond to normalized numbers of mapped reads per million reads per 200 bp bin.

#### Calculation of read coverage

In wild-type and *H3^K36A and R^* mutant conditions, H3K36me2 and H3K27me3 ChIP-seq read coverages across gene bodies were computed on genomic intervals starting 750 bp upstream transcription start sites and ending 750 bp downstream transcription termination sites. Read coverage is defined as the normalized number of mapped reads per million reads from a ChIP-seq dataset divided by the number of mapped reads per million reads from the corresponding input dataset across a genomic region. Among the *D. melanogaster* Refseq genes, approximately 10800 and 9200 are overlapping with H3K36me2 enriched regions, approximately 1030 and 1030 genes are overlapping with high-level H3K27me3 domains and 5400 and 6300 are localized in other genomic regions in embryos and larvae, respectively.

### *Drosophila* nuclear and cell extracts for western blot analysis

For embryonic total nuclear extracts, nuclei from 21-24 hr old *wt*, *H3^K36A^* or *H3^K36A^* mutant embryos were purified and quantified as described (Bonnet et al., 2019). Pellets of nuclei were resuspended in appropriate volumes of SDS sample buffer proportional to the number of nuclei in each pellet. Extracts were then sonicated in a Bioruptor instrument (Diagenode) (8 cycles (30 sec ON / 30 sec OFF), high power mode), incubated at 75° C for 5 min and insoluble material was removed by centrifugation at 14000 rpm for 1 mn at r.t..

Total cell extracts from imaginal disc tissues were prepared by resuspending hand-dissected disc tissues in SDS sample buffer. Extracts were then sonicated, incubated at 75°C for 5 min and insoluble material was removed by centrifugation.

### Antibodies

For ChIP analysis:

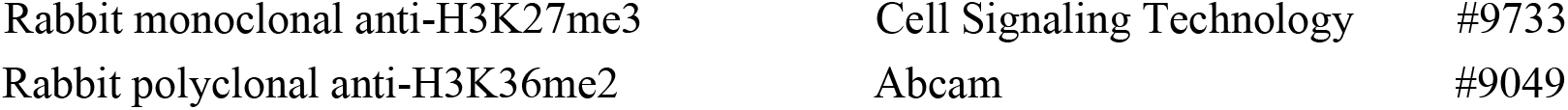

For Western blot analysis on embryonic and larval extracts:

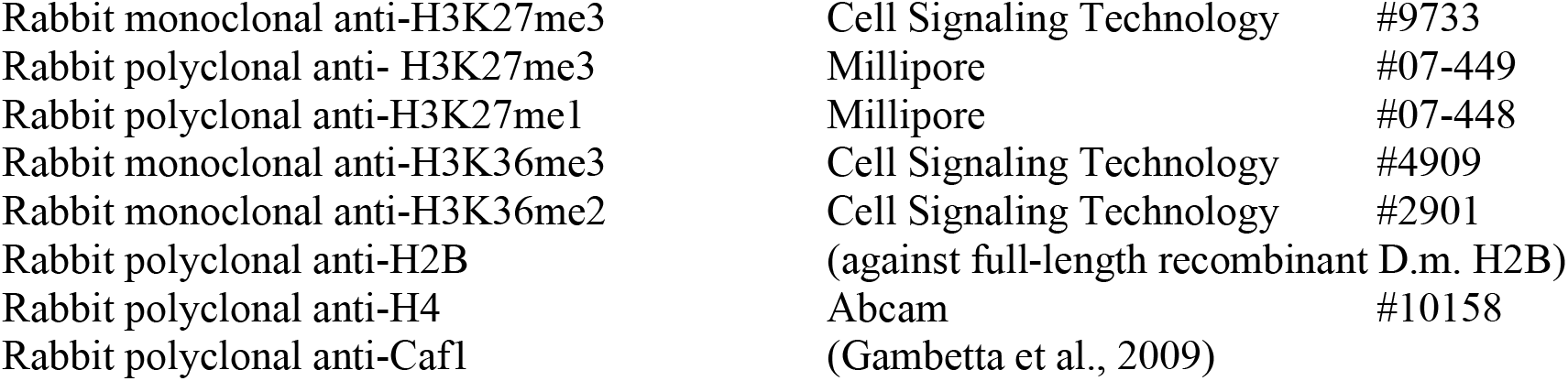

For immunohistochemistry and immunofluorescence analysis:

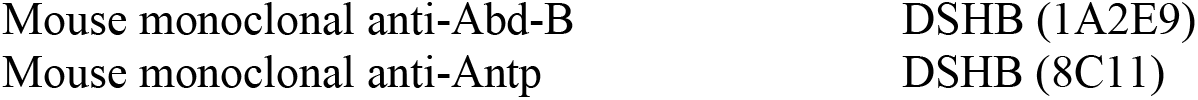

## ACKNOWLEDGMENTS

We thank Eva Nogales for generous advice, sharing of expertise and for hosting K.F. for grid preparation. We thank Tom Cech for stimulating discussions. We thank J.R. Prabu for excellent computing support, S.Uebel, E.Weyher, R.Kim and A.Yeroslaviz of the MPIB core facilities for excellent technical support and S.Schkoelziger and S.Schmähling for help with some of the experiments. This work was supported by the Deutsche Forschungsgemeinschaft (SFB1064) and the MPG. ChIP-seq data have been deposited in GEO (accession number: GSE148254).

---

**SUPPLEMENTARY FIGURE 1.**
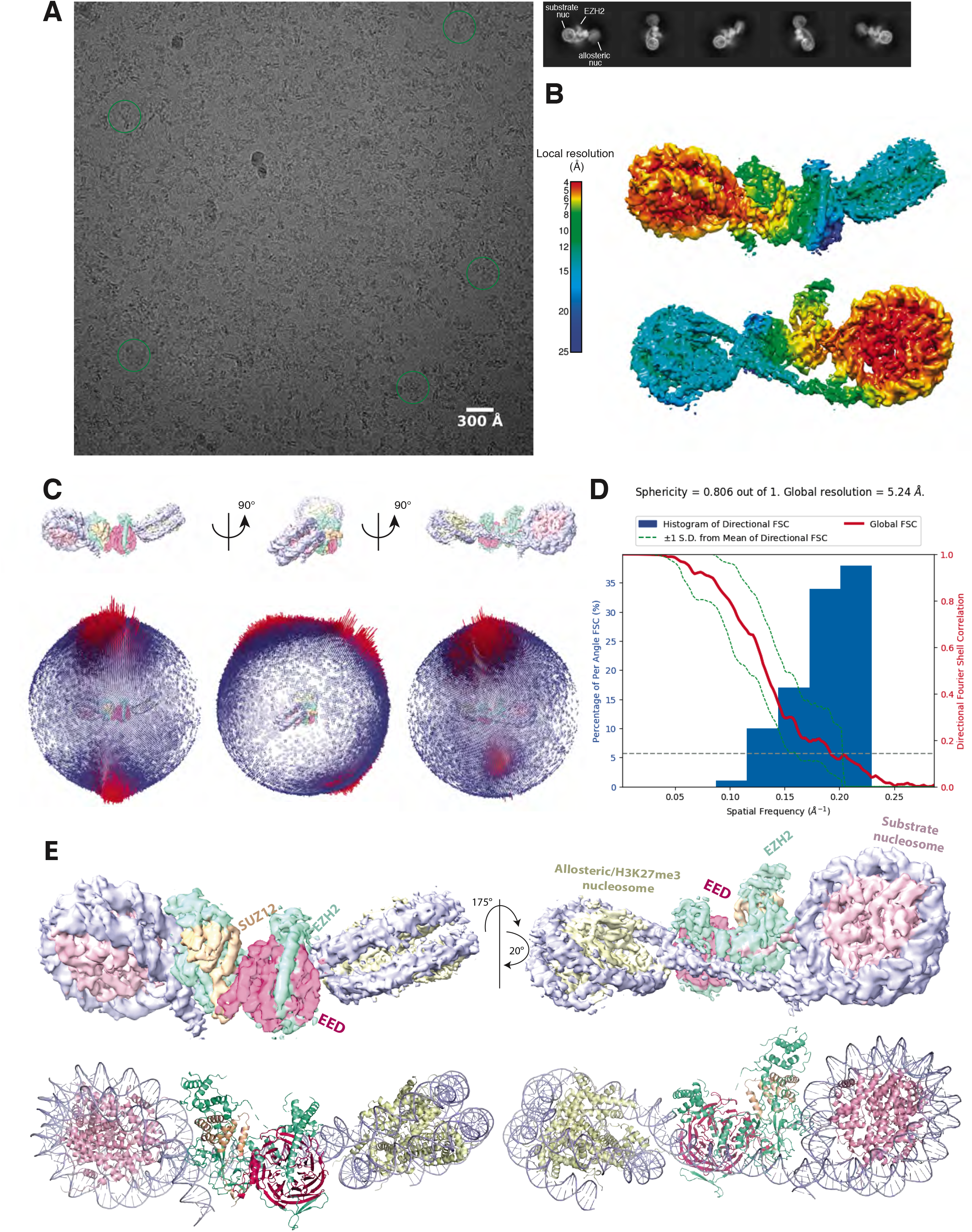
Initial Cryo-EM analysis of the PHF1-PRC2:di-Nuc complex (related to Fig. 1). (**A**) Representative micrograph of the cryo-EM dataset (left) and reference-free 2D classes from particles picked without templates (right) (performed to ensure that no bias was introduced through templates picking and references in 3D classification).. Circles indicate particles, which were picked with templates and directly subjected to 3D analysis (see **fig. S2**). (**B**) Local resolution estimation of the 5.2 Å overall PHF1-PRC2:di-Nuc map. The substrate nucleosome and the adjacent part of EZH2 are well resolved (colors red to yellow). (**C**) Spherical angular distribution of particles included in the final reconstruction of PHF1-PRC2:di-Nuc. (**D**) Output from the 3DFSC Processing Server (https://3dfsc.salk.edu/ (Tan et al., 2017)) showing the Fourier Shell Correlation (FSC) as a function of spatial frequency, generated from masked independent half maps of PRC2:diNuc: global FSC (red), directional FSC (blue histogram) and deviation from mean (spread, green dotted line). The nominal overall resolution of 5.24 Å was estimated according to the gold standard FSC cutoff of 0.143 (grey dotted line) (Rosenthal and Henderson, 2003). Sphericity is an indication for anisotropy and amounts to 0.806 in this data. The minor directional anisotropy of the data can be explained by the slightly preferred orientation and missing views as seen in (**C**). (**E**) Top: Refined and postprocessed cryo-EM density map of overall PHF1-PRC2:di-Nuc colored according to the subunit organization. Bottom: pseudoatomic model of fitted crystal structure of the human PRC2 catalytic lobe (PDB: 5HYN) and a di-Nuc model with 35 bp linker DNA (Poepsel et al., 2018), including PDB 1AOI.

**SUPPLEMENTARY FIGURE 2.**
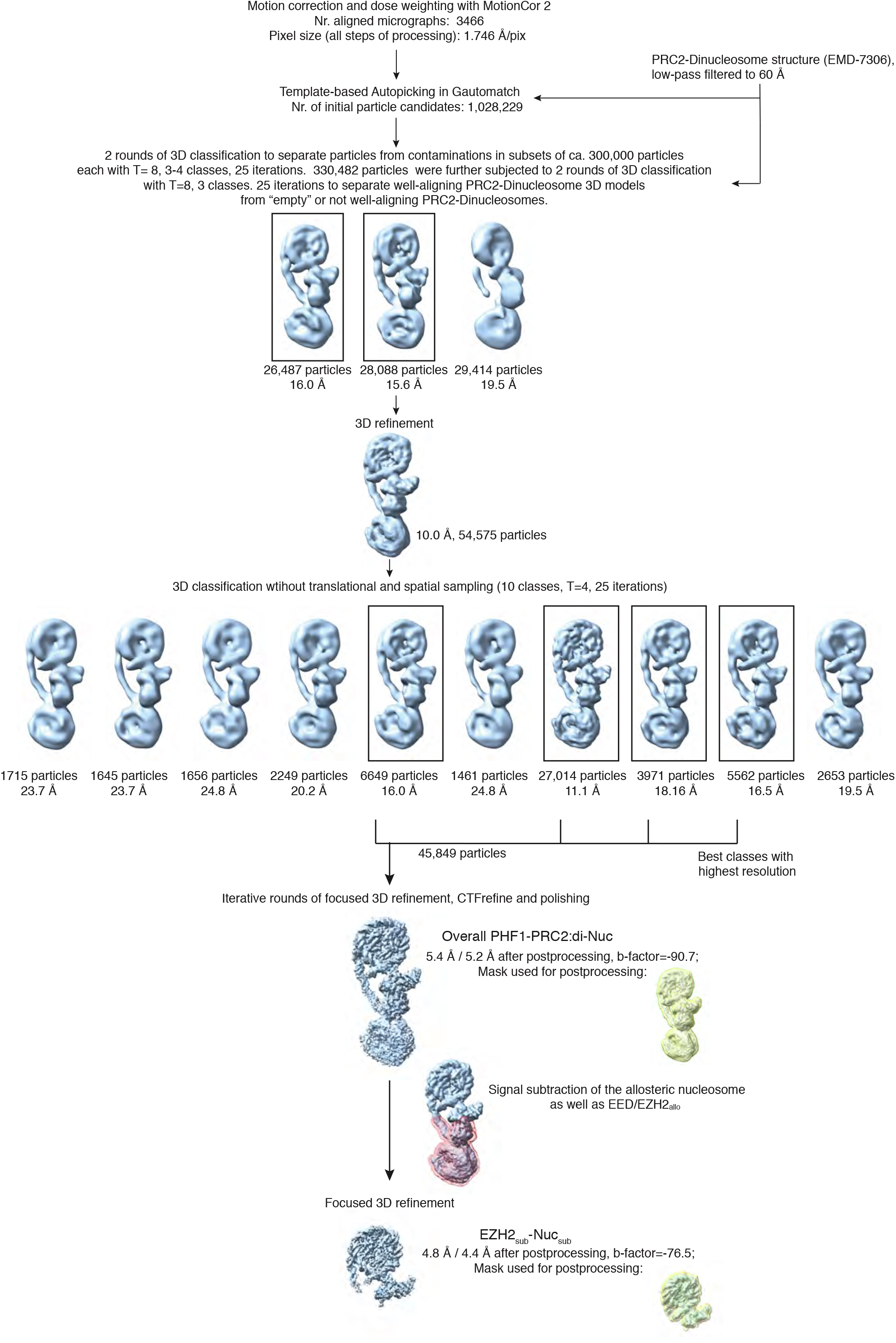
Overview of the cryo-EM Data-Processing and Particle Sorting Scheme (related to Figure 1). Processing and particle sorting scheme, also described in Methods. Squares indicate 3D classes (and corresponding particles) chosen for further processing steps based on their nominal global resolution values, translational and rotational accuracy and the presence of detailed structural information. Two final reconstructions were obtained in this study: Overall PHF1-PRC2:di-Nuc, and EZH2_sub_-Nuc_sub_ after performing signal subtraction (mask indicated in pink) and focused refinement. Masks used for postprocessing are shown in yellow.

**SUPPLEMENTARY FIGURE 3.**
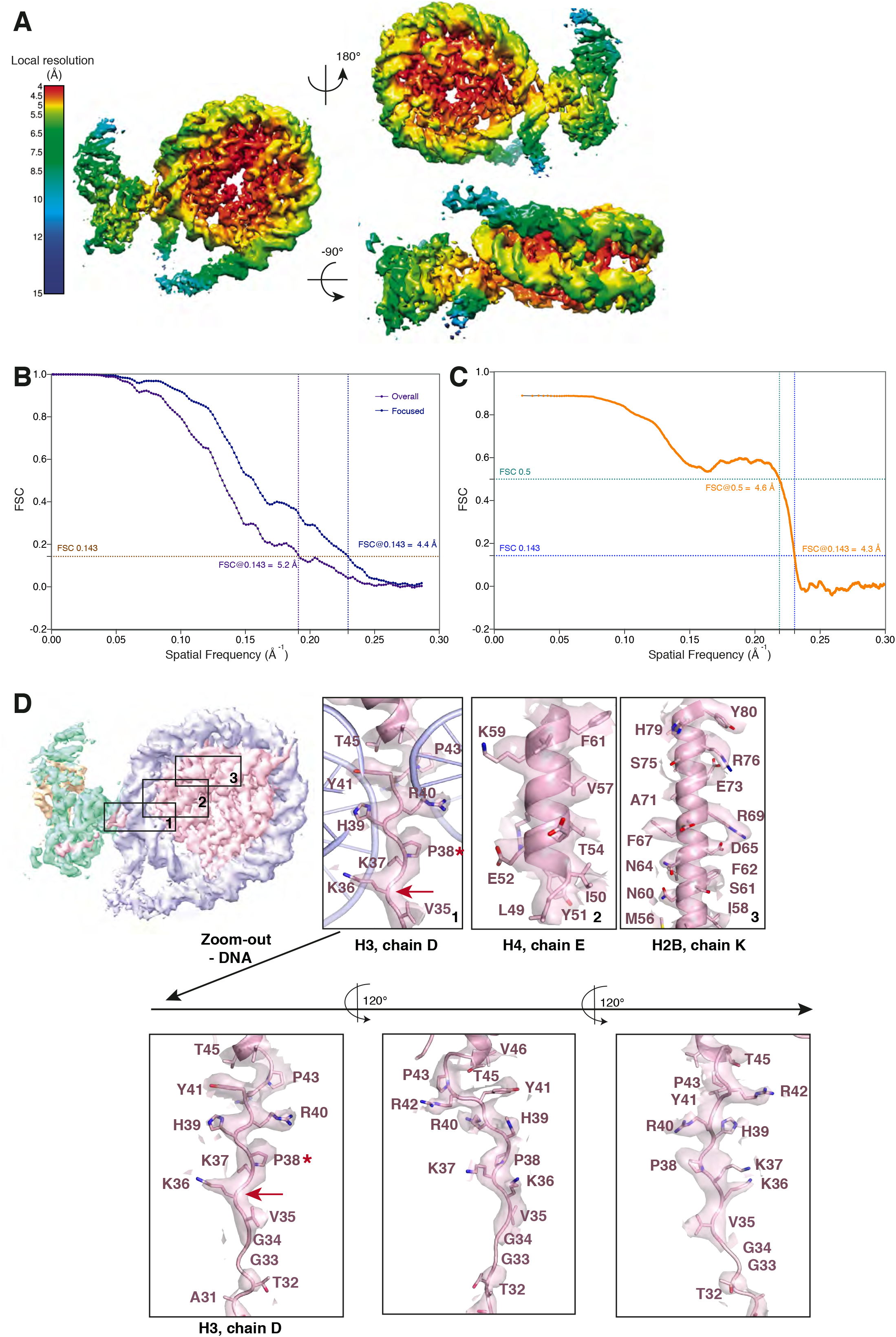
Cryo-EM analysis of the focused EZH2_sub_-Nuc_sub_ map (related to Fig. 1). (**A**) Local resolution estimation of the focused 4.4 Å EZH2_sub_:Nuc_sub_ reconstruction. Regions in the nucleosome core as well as the adjacent regions including parts of the H3 N-terminus close to the exit side of the nucleosome are well resolved (4.0 – 5.5 Å). Regions close to the mask, especially the nucleosomal DNA and parts of EZH2, are less well resolved (colors green to blue). (**B**) Global FSC generated from masked independent half maps of EZH2_sub_-Nuc_sub_ (Focused, blue line) and the overall PHF1-PRC2:di-Nuc (Overall, violet line) were plotted against spatial frequency. The resolution of 4.4 Å for for EZH2_sub_-Nuc_sub_ map and 5.2 Å for the overall PHF1-PRC2:di-Nuc map were estimated according to the gold standard FSC cutoff of 0.143 (brown dotted line) (Rosenthal and Henderson, 2003) (**C**) FSC between the atomic model and the masked (applied in Phenix) map of EZH2_sub_-Nuc_sub_ after real-space refinement (Afonine et al., 2018). Green line represents the cut-off at 0.5 (4.6 Å) and blue line represents the cut-off at 0.143 (4.3 Å) (see also Table 1) (Henderson et al., 2012; Rosenthal and Henderson, 2003; Rosenthal and Rubinstein, 2015). (**D**) Selected regions within EZH2_sub_-Nuc_sub_ showing side chain density, e.g. K36 (red arrow). A red asterisk indicates the last residue of the H3 tail visible in known crystal structures (usually P38 or H39). The quality of the map around K36 is shown as a separate zoom-out below and in three different views to demonstrate the lack of anisotropy present in the density.

**SUPPLEMENTARY FIGURE 4.**
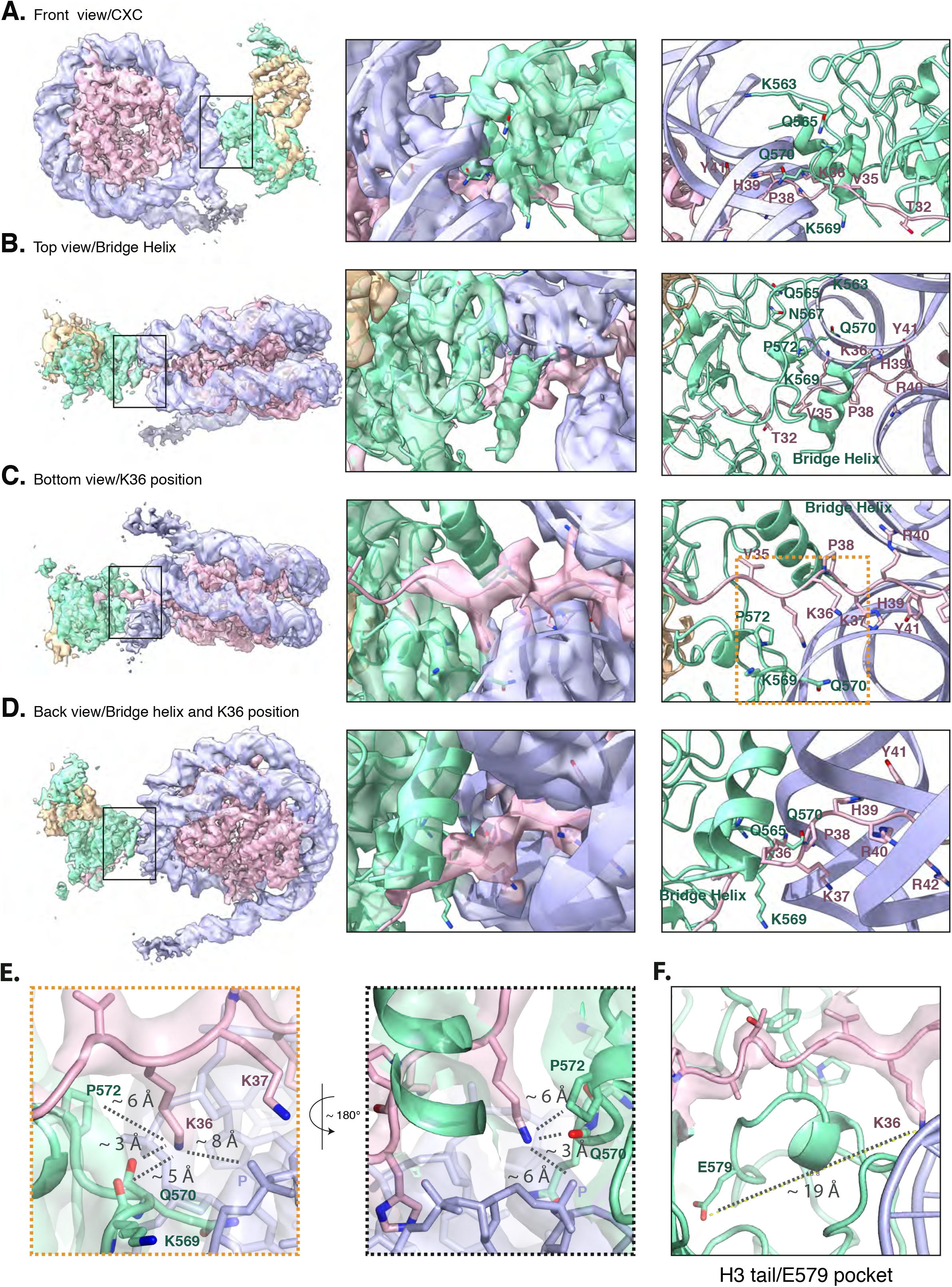
The improved map of the interaction between EZH2 and the substrate nucleosome after focused refinement reveals location of H3K36 and it’s environment (related to Fig. 1). (**A**) The front view of EZH2_sub_-Nuc_sub_ cryo-EM density and model shows details of the EZH2_CXC_ interaction with nucleosomal DNA. (**B**) The top view of EZH2_sub_-Nuc_sub_ cryo-EM shows a tubular density into which based on recent findings of (Kasinath et al., 2020) a helix was built. The bridge helix, which based on this study is likely constituted of the EZH2 residues 497-511, is located above V35 of the H3 tail. As can be seen when observing the density-modified map (Terwilliger et al., 2019) of EZH2_sub_-Nuc_sub_ at lower threshold, it presumably engages in interactions with the nucleosomal DNA, the H3 tail and EZH2, as described in greater detail in (Kasinath et al., 2020). (**C**) The bottom view of EZH2_sub_-Nuc_sub_ cryo-EM density and model shows details of the vicinity of K36 with the corresponding density for the H3 tail, EZH2 and nucleosomal DNA. The orange square indicates the region shown as a zoom-in in (**E**). (**D**) The back view of EZH2_sub_-Nuc_sub_ cryo-EM density and model shows details of the location of K36 and the bridge helix. (**E**) Zoom-in views of H3K36 and it’s chemical environment. Approximate distances of the epsilon-amino group of H3K36 to the nearest residues are indicated with a dotted grey line. (**F**) Location of the Glu-579 pocket (Jani et al., 2019) in the EZH2_sub_:Nuc_sub_ reconstruction and it’s distance to H3K36 (app. 19 Å). The described mechanism by Jani et al (Jani et al., 2019) involving recognition of H3K36 by Glu-579 is incompatible with the presented structural data as the location differs significantly and major rearrangements as the relocation of the helix-loop region between residues 564-576 would be necessary to avoid the given steric and geometric hindrance and allow for potential interaction.

**SUPPLEMENTARY FIGURE 5.**
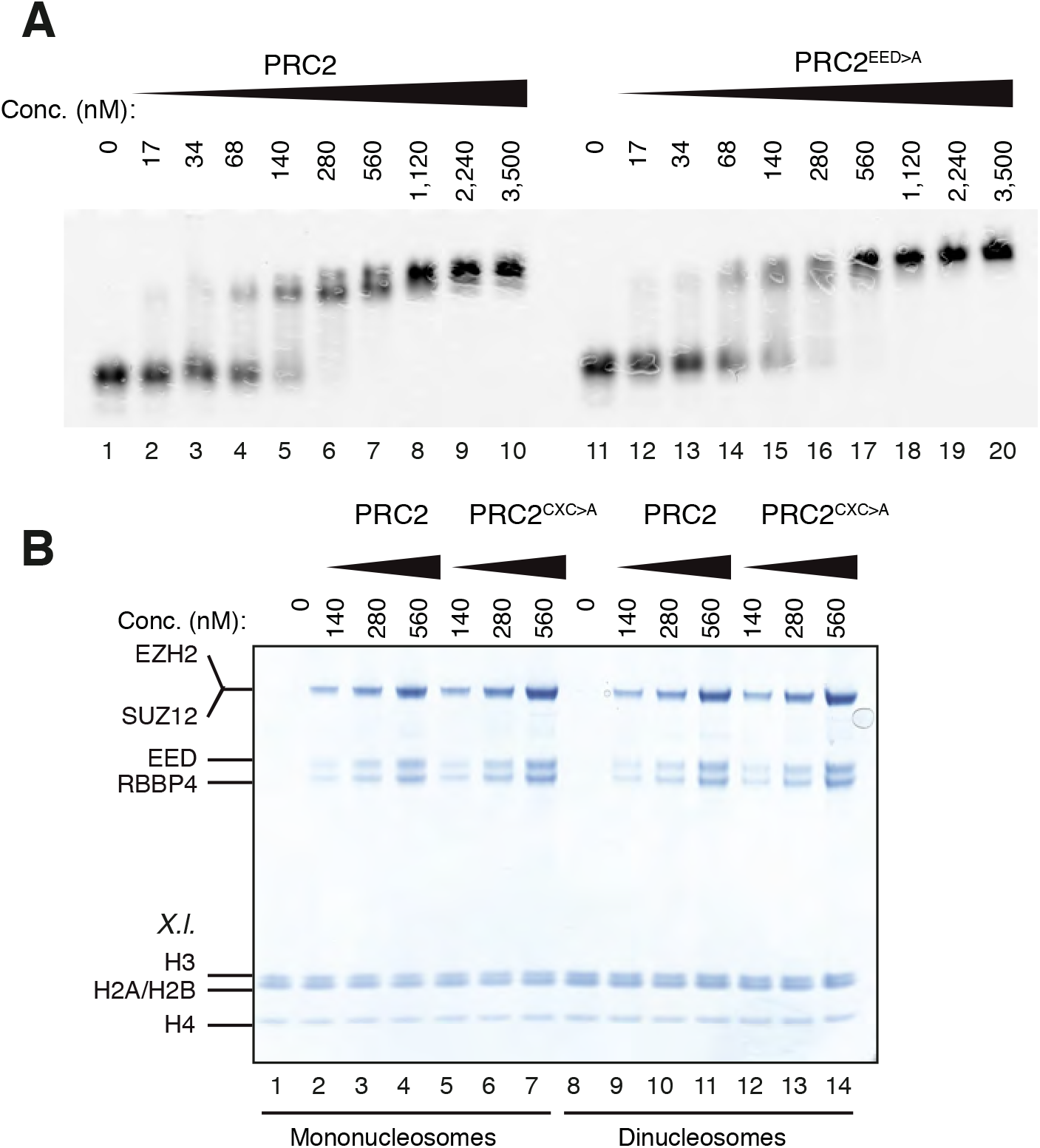
The EZH2_CXC_-DNA interaction interface is critical for H3K27 methylation on nucleosomes (related to Fig. 2). (**A**) Binding reactions with indicated concentrations of PRC2 (lanes 1-10) or PRC2^EED>A^ (lanes 11-20) and 45 nM 6-carboxyfluorescein-labeled mononucleosomes, analyzed by EMSA on 1.2% agarose gels. (**B**) Coomassie-stained 4-12% SDS-PAGE of the HMTase reactions shown in **Fig. 2C**. *Xenopus laevis (X.l.*) nucleosomes were used for these experiments. The short 5-kDa PHF_C_ fragment is not visible on this gel.

**SUPPLEMENTARY FIGURE 6.**
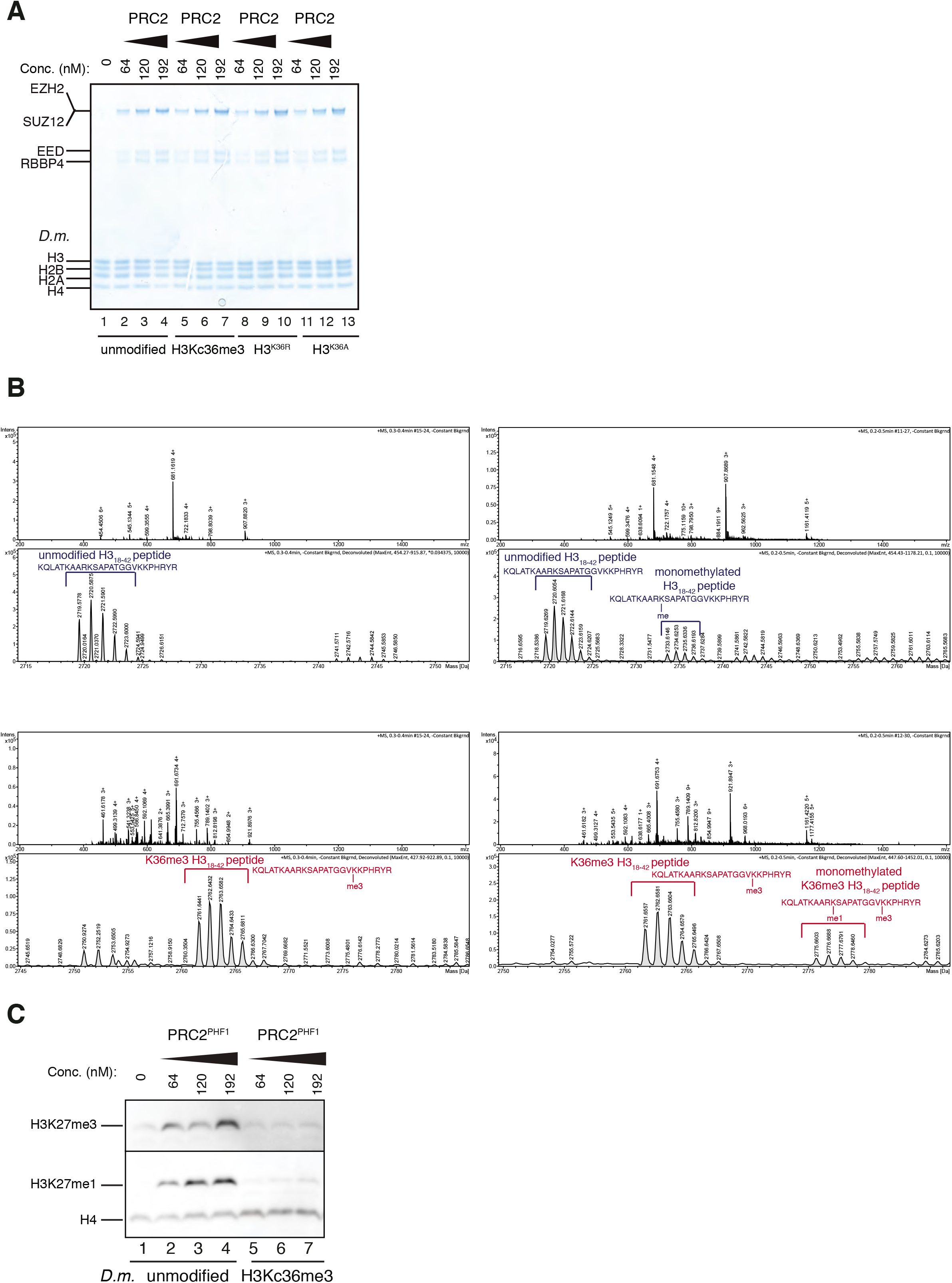
Accommodation of unmodified H3K36 in the EZH2_CXC_-DNA interaction interface is essential for H3K27 methylation on nucleosomes and PHF1-PRC2 (related to Fig. 3). (**A**) Coomassie-stained 4-12% SDS-PAGE of the HMTase reactions shown in **Fig. 3C**. *Drosophila melanogaster (D.m.*) nucleosomes were used for these experiments. The short 5-kDa PHF_C_ fragment is not visible on this gel. (**B**) Full ESI MS spectra (upper part) and full deconvoluted MS spectra (lower part) shown for input peptides without PRC2 as a control (left) and with PRC2 (right) to ensure no overlapping between possible adduct peaks and monomethylation peaks. (**C**) Western Blot (WB) analysis of HMTase reactions with full-length PHF1-PRC2 on unmodified (lanes 1-4) or H3Kc36me3 (lanes 5-7) mononucleosomes (446 nM).

**SUPPLEMENTARY FIGURE 7.**
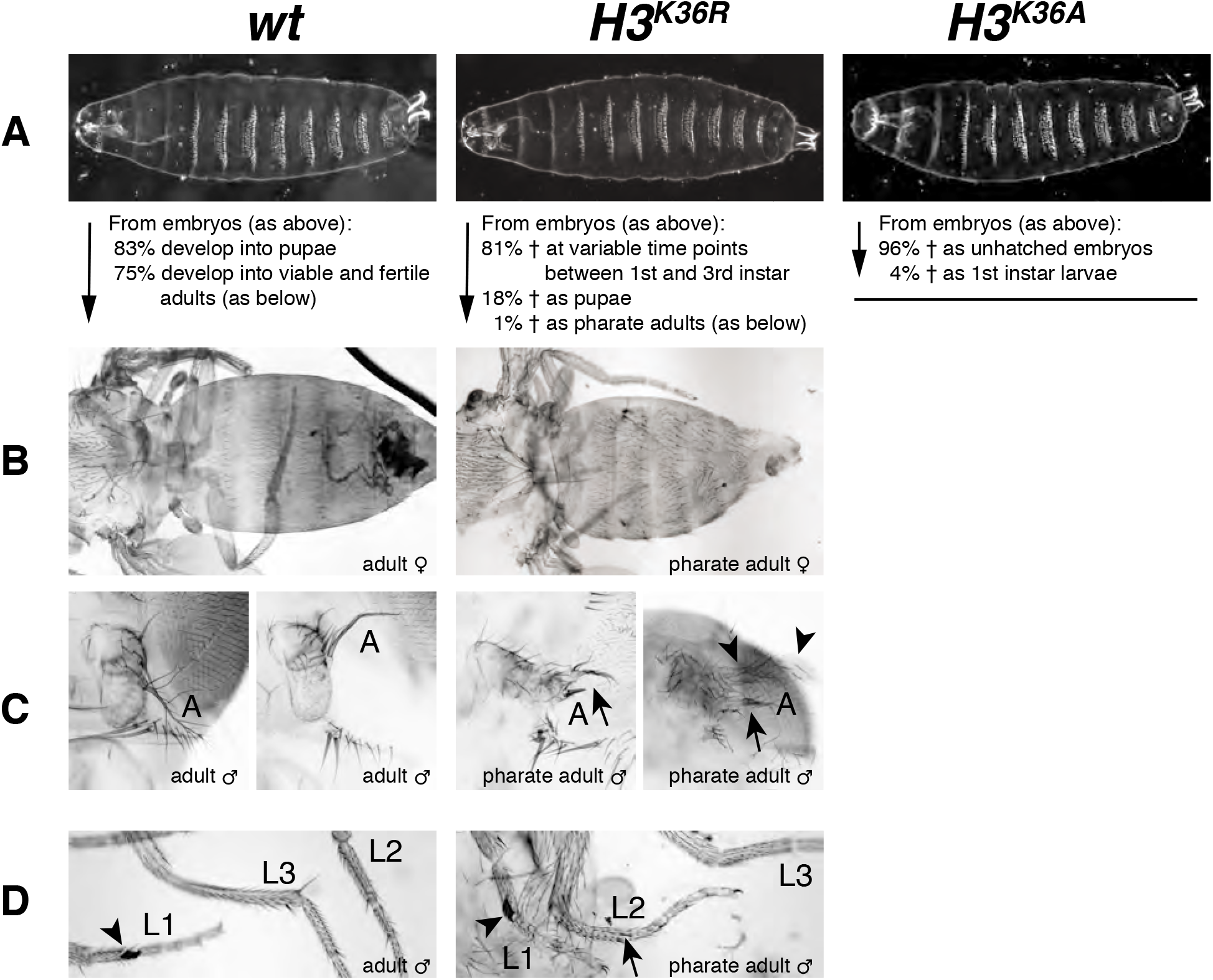
*Drosophila* with H3^K36R^ or H3^K36A^ mutant chromatin arrest development at different stages. (**A**) Ventral views of cuticles from wildtype (*wt*), *H3^K36R^*, or *H3^K36A^* mutant embryos. Note that the cuticle pattern of the mutant animals is indistinguishable from that of the *wt* embryo. Below: for each genotype, the fraction of embryos that developed into larve, pupae, pharate adults or viable adults is listed. The fraction was determined by monitoring the development of collected hatched 1^st^ instar larvae (*wt*: *n*=300, *H3^K36R^: n*=2000) or unhatched embryos (*H3^K36A^: n*=200). The GFP marker on the Balancer chromosomes was used for identifying *H3^K36R^* and or *H3^K36A^* mutants. (**B**) Dorsal views of the posterior portion of the thorax and of the abdomen. From 2000 hatched *H3^K36R^* mutant 1^st^ instar larvae, a total of 18 pharate adults was recovered. Most *H3^K36R^* mutant pharate adults showed a relatively normal overall body patterning apart from the homeotic transformations illustrated below. (**C**) Frontal view of adult heads illustrating the antenna-to-leg transformation in *H3^K36R^* mutant pharate adults. The antenna-to-leg transformation in *H3^K36R^* mutant animals ranged from mild (arrows) to more extensive transformations with formation of leg-like structures such as in this extreme case (arrowheads). (**D**) The sex comb in males is normally only present on the protoracic (L1) legs (arrowheads). Among the *H3^K36R^* mutant pharate adult males recovered (*n*=13), five showed one or several extra sex comb teeth (arrow) on the meso- (L2) or metathoracic (L3) legs. Extra sex comb teeth in adults are a hallmark phenotype of Polycomb mutants.

**SUPPLEMENTARY FIGURE 8.**
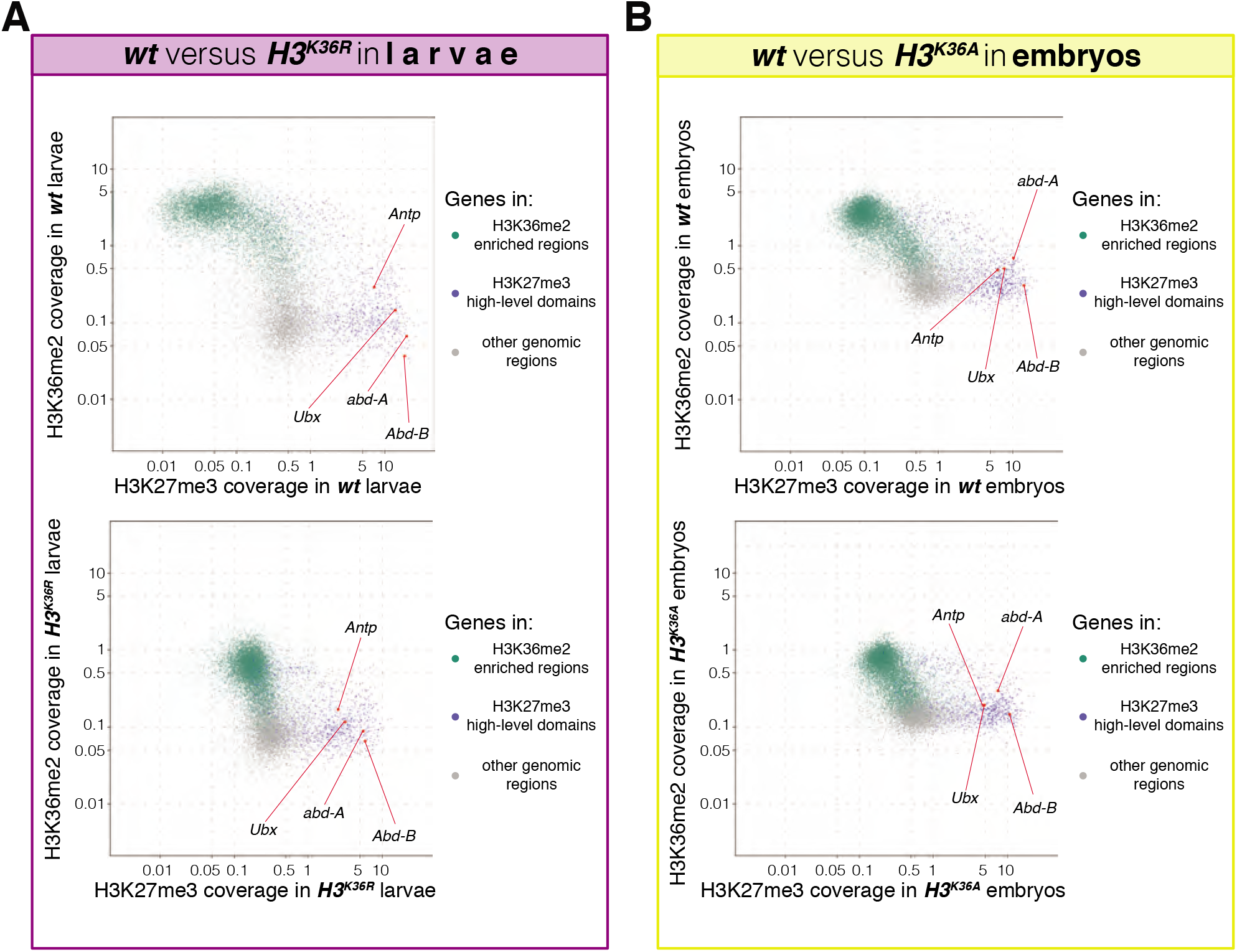
*H3^K36R^* and *H3^K36A^* mutants show altered H3K36me2 and H3K27me3 profiles (related to Fig. 4) (**A**) Top, scatter plots showing H3K36me2 coverage in relation to H3K27me3 coverage in *wt* larvae. Green dots represent 9200 gene bodies overlapping with genomic intervals showing H3K36me2 enrichment, blue dots represent 1030 gene bodies overlapping with genomic intervals defined as high-level H3K27me3 domains (Bonnet et al., 2019), and grey dots represent 6300 gene bodies showing no enrichment for either methylation mark in larvae (see Materials and Methods). Bottom, scatter plot showing the H3K36me2 read coverage in relation to H3K27me3 read coverage in *H3^K36R^* mutant larvae. (**B**) As in (**A**) but showing H3K36me2 coverage in relation to H3K27me3 coverage in *wt* embryos (top) and in *H3^K36A^* mutant embryos (bottom). Green dots represent 10800 gene bodies overlapping with genomic intervals showing H3K36me2 enrichment, blue dots represent 1030 gene bodies overlapping with genomic intervals defined as high-level H3K27me3 domains (Bonnet et al., 2019), and grey dots represent 5400 gene bodies showing no enrichment for either methylation mark in embryos (see Materials and Methods).

**Table S1.** Cryo electron microscopy data collection summary, processing statistics and model.

| Cryo electron microscopy data collection (applicable to Overall PHF1-PRC2:diNuc and EZH2<sub>sub</sub>-Nuc<sub>sub</sub> maps) |  |  |
| --- | --- | --- |
| Microscope | FEI Titan Krios GII |  |
| Voltage (kV) | 300 |  |
| Camera | Gatan K2-Summit |  |
| Energy Filter | Gatan Quantum-LS (GIF) |  |
| Pixel size (Å/pix) (calibrated) | 1.75 |  |
| Nominal magnification (x) | 81000 |  |
| Preset target global defocus range (µm) | 0.5 - 3.5 |  |
| Total electron exposure (fluence, e<sup>-</sup>/Å<sup>2</sup>) | 52,96 |  |
| Exposure rate (flux) (e<sup>-</sup>/ Å<sup>2</sup>/s) | 3,47 |  |
| Nr. of frames collected per micrograph | 60 |  |
| Energy filter slit width (eV) | 20 |  |
| Automation Software | SerialEM |  |
| 3D reconstruction (applicable to Overall PHF1-PRC2:diNuc and EZH2<sub>sub</sub>-Nuc<sub>sub</sub> maps) |  |  |
| Number of movies | 3466 |  |
| Initially selected particle candidates | 1,028,229 |  |
| Final number of particles | 45,849 |  |
|  | Overall PHF1-PRC2:diNuc | EZH2<sub>sub</sub>-Nuc<sub>sub</sub> |
| Resolution FSC independent halfmaps (0.143) masked (Å)<sup>a</sup> | 5.24 | 4.36 |
| Local resolution range (Å) | 4.01 - 24.97 | 4.01 – 15.00 |
| Sharpening B-factor (Å<sup>2</sup>) | -90.7 | -76.5 |
| Refinement |  |  |
| EZH2<sub>sub</sub>-Nuc<sub>sub</sub> |  |  |
| No. atoms |  | 28866 (Hydrogens: 13086) |
| Residues |  | Protein: 1171 Nucleotide: 312 |
| Ligands |  | ZN: 8 |
| CC<sub>mask</sub>, CC<sub>box</sub>, CC<sub>peaks</sub>, CC<sub>volume</sub><sup>b</sup> |  | 0.75, 0.83, 0.66, 0.74 |
| Mean CC for ligands |  | 0.72 |
| ResolutionFSC masked map vs. model (0/0.143/0.5) (Å)<sup>b</sup> |  | 4.3/4.3/4.6 |
| R.m.s. deviations |  |  |
| Bond lengths (Å) |  | 0.004 |
| Bond angles (°) |  | 0.642 |
| Ramachandran favored (%) |  | 97.51 |
| Ramachandran gen. allowed (%) |  | 2.32 |
| Ramachandran disallowed (%) |  | 0.17 |
| MolProbity score |  | 1.53 |
| Clash score |  | 7.97 |
| ADP (B factors) |  |  |
| Iso/Aniso (#) |  | 15925/0 |
| Min/max/mean |  |  |
| Protein |  | 65.89/264.21/121.82 |
| Nucleotide |  | 97.31/221.62/132.65 |
| Ligand |  | 151.54/251.77/201.51 |
| Rotamer outliers (%) |  | 0.00 |
| Cβ outliers (%) | | 0.00 |
| CaBLAM outliers (%) |  | 1.59 |

<sup>a</sup>according to the Fourier Shell Correlation (FSC) cut-off criterion of 0.143 defined in (Rosenthal and Henderson, 2003)
<sup>b</sup>according to the map-vs.-model Correlation Coefficient definitions in (Afonine et al., 2018a)

**Table S2.**

**Number of aligned reads to the *D. melanogaster* and *D. pseudoobscura* genomes from ChIP and input datasets and normalization process (related to Figure 4 and Supplementary Figure 8).**

**Movie S1**

**Cryo-EM structure of the PHF1-PRC2:di-Nuc complex (related to Figure 1).**

